# Nuclei function as wedges to pry open spaces and promote collective, confined cell migration *in vivo*

**DOI:** 10.1101/2021.12.16.473064

**Authors:** Lauren Penfield, Denise J. Montell

## Abstract

Cells migrate collectively through confined environments during development and cancer metastasis. The nucleus, a stiff organelle, impedes single cells from squeezing into narrow channels within artificial environments, but how nuclei affect collective cell migration into compact tissues *in vivo* is unknown. Here, we use border cells in the fly ovary to study nuclear dynamics in collective, confined *in vivo* migration. Border cells delaminate from the follicular epithelium and squeeze into tiny spaces between cells called nurse cells. The lead cell nucleus transiently deforms as it extends into the lead cell protrusion, which then widens. The nuclei of follower cells deform less. Depletion of the *Drosophila* B-type lamin, Lam, compromises nuclear integrity, hinders expansion of a leading protrusion, and impedes border cell movement. Cortical myosin II accumulates behind the nucleus and moves it into the protrusion in wildtype but not Lam-depleted cells. These data suggest that the nucleus stabilizes lead cell protrusions to help cells wedge open spaces between nurse cells.

**Summary:** During *Drosophila* border cell migration, the B-type lamin Lam maintains nuclear envelope integrity, stabilizes the lead cell protrusion, and promotes cluster invasion between nurse cells. The nucleus may function as a wedge to prom collective, confined *in vivo* movement.

## Introduction

Collective cell migration is critical for development and wound healing, and promotes cancer metastasis (Rørth, 2009; Friedl and Gilmour, 2009). One challenge for collective groups moving *in vivo* is the difficulty of squeezing into small spaces between cells, for example during intravasation and extravasation (Stoletov et al., 2010; Wyckoff et al., 2000; Reymond et al., 2013), or through dense extracellular matrix (ECM). How groups of cells physically crawl into narrow paths and navigate complex tissue environments is not well understood.

Nuclear deformations have been widely observed in cells migrating through tight spaces *in vivo* (Stoletov et al., 2010; Yamauchi et al., 2005; Norden et al., 2009; Denais et al., 2016; Raab et al., 2016; Kalukula et al., 2022). Since the nucleus can pose a physical barrier to movement through confined spaces (Calero-Cuenca et al., 2018; Friedl et al., 2011), a major research focus is to determine how nuclei contribute to or obstruct confined migrations through natural environments. Many studies have focused on the nucleus in confined, single cell migration (Yamada and Sixt, 2019; McGregor et al., 2016), while how nuclei affect collective, confined *in vivo* migration is less understood.

The nucleus is typically the largest and stiffest organelle and is mechanically supported by networks of lamin filaments. A- and B-type lamins are proteins that assemble into distinct intermediate filament networks underneath the nuclear envelope (NE), and are primary contributors to nuclear mechanics (Hetzer, 2010; Davidson and Lammerding, 2014; Wintner et al., 2020; Lammerding et al., 2004). Higher expression of lamins, particularly A-type lamins, increases nuclear stiffness (Swift et al., 2013; Ferrera et al., 2014). A-type lamin tends to be expressed at higher levels in stiff tissues while B-type expression is higher during development and in soft tissues such as the brain (Swift et al., 2013; Hutchison, 2014). *In vitro*, mouse embryonic fibroblasts, cancer cell lines, mesenchymal stem cells, and hematopoietic cells depleted of A-type lamins have more deformable nuclei and migrate faster through narrow artificial channels or dense ECM (Rowat et al., 2013; Davidson et al., 2014; Harada et al., 2014; Shin et al., 2013), so nuclear stiffness can impede confined migration. In support of this idea, fast migratory cells, such as circulating white blood cells, downregulate A-type lamins (Shin et al., 2013) and are able to move rapidly through narrow channels in silicone devices (Rowat et al., 2013; Raab et al., 2016; Thiam et al., 2016). In contrast to the notion that nuclear rigidity is a barrier to migration, several studies report that lamin A/C depletion reduces the ability of multiple cell types including mesenchymal stem cells, melanoma cells, and dendritic cells, to move in confinement (Lee et al., 2021; Lomakin et al., 2020). Thus, lamins may have distinct effects on migration depending on the cell type, migration environment, and migration mode.

There are multiple, not necessarily mutually exclusive, models for how nuclear mechanical properties contribute to confined migration. When the nuclei of a variety of cell types including HeLa or primary zebrafish progenitor cells are compressed, they respond by unfolding indentations in the nuclear envelope and triggering phospholipase A_2_ (PLA_2_)- stimulated cortical actomyosin contractions, which promote extensive blebbing of the plasma membrane and escape from the confinement (Lomakin et al., 2020; Venturini et al., 2020). This mechanism, which is activated when the cell is confined to a space smaller than the nuclear diameter, is referred to as the nuclear ruler (Lomakin et al., 2020; Venturini et al., 2020). The nuclear piston model, initially reported in primary human fibroblasts (Petrie et al., 2014) and later observed other cell types including confined, mesenchymal stem cells (Lee et al., 2021), proposes that hydrostatic pressure builds up in confined cells, activating mechanosensitive ion channels that open and cause swelling of large, “lobopodial” protrusions (Lee et al., 2021). Again, blebbing accompanies the response to confinement. In leukocytes, the nucleus is positioned near the front of the cell and acts as a mechanical gauge, allowing the cell to identify the path of least resistance (Renkawitz et al., 2019). A key open question is how nuclei sense, respond, and contribute to confined cell migration *in vivo*, particularly in cells that migrate collectively in between other cells.

Here, we use the border cell cluster in the *Drosophila* ovary as a model to study how nuclei change shape during and contribute to collective, confined, *in vivo* cell migration. We show that nuclei rapidly deform as cell clusters move in between tightly apposed nurse cells. Lead cell nuclei undergo the most significant shape changes, elongating as they move into forward-directed protrusions, and then recovering elastically. Nuclear movement correlates with protrusion expansion. We further show that reduced lamin expression delays migration. While both A- and B-type lamins are expressed, only the B-type lamin, Lam, is required to maintain nuclear integrity and promote border cell movement in between nurse cells. Lam-depleted cells extend transient forward protrusions that do not enlarge properly, as well as ectopic protrusions, ultimately resulting in uncoordinated movement and failed invasion between nurse cells. Cortical nonmuscle myosin II flashes move the nucleus into protrusions in control cells, while the flashes move past Lam-depleted nuclei and instead constrict the protrusion. We did not detect hallmarks of the nuclear piston or nuclear ruler, such as blebbing, in border cells. The data suggest that, rather than a ruler, gauge, or piston, the nucleus promotes invasion of the border cell cluster into a space that is initially much smaller than even a single nucleus, possibly by functioning as a wedge.

## Results

### Leader cell nuclei transiently deform as the lead cell protrusion widens and border cells move between nurse cells

To study the role of nuclei in collective cell migration, we used the well-established model of border cell migration in the *Drosophila* egg chamber. At stage 9, the border cell cluster delaminates from a layer of epithelial cells and squeezes in between germline cells, termed nurse cells, to reach the oocyte by stage 10 (Fig 1A-C, (Montell et al., 2012)). Border cells consist of a pair of inner, nonmotile polar cells and 4-6 outer, motile cells (Fig 1B). Typically, one outer border cell extends a large protrusion toward the oocyte and leads the cluster, though the leader can change over time. Lead border cells extend and retract protrusions, probing for chemoattractants and available space. The border cells select the central path between the nurse cells because it contains slightly larger spaces (Dai et al., 2020) (Fig. 1D); however even the largest spaces are much smaller than the cluster.

**Figure 1.**
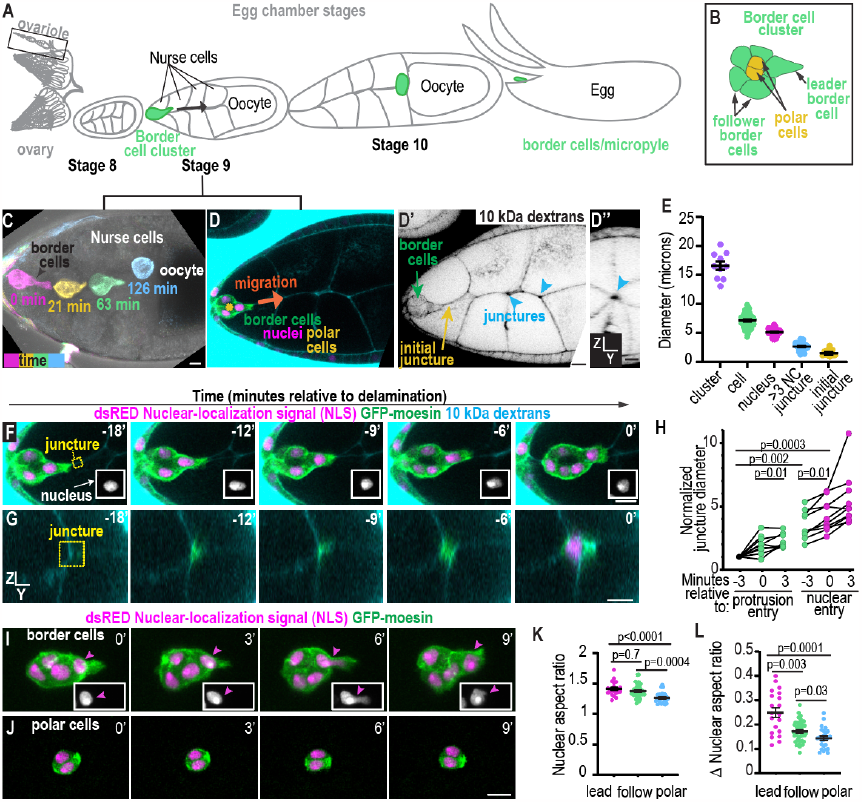
Leader cell nuclei transiently deform during the onset of border cell migration. A. Overview of the *Drosophila* ovary, ovariole, and egg chamber stages. B. Illustration of the border cells (green) and polar cells (purple). C. A max projection of images from a timelapse of a stage 9 egg chamber to show the border cell cluster movement over time. The egg chamber is shown with Differential Interference Contrast (DIC) and border cells expressing GFP-moesin are labeled with a different color for each timepoint. D. A stage 9 egg chamber expressing fruitlessGal4;UAS-GFP-moesin (border cells);UASdsRED.Nuclear localization signal (NLS) (nuclei) and incubated with 10 kDa dextran-Alexa647 to label junctures. D’-D’’ show the inverted fluorescent image of the 10 kDa dextrans in both XY and YZ views to show possible paths for border cells to migrate on. Arrowheads mark central extracellular spaces which the border cells select to migrate in. Scale bars: 10 ***μ***m E. Individual and mean +/-standard error of the mean (SEM) values for diameters of the indicated features. A one way ANOVA followed by a post-hoc Tukey was performed, all diameters are significantly different from one another, p<0.0001 for all comparisons except initial juncture versus >3 NC juncture, p =0.004. Measurements from 10 egg chambers. F-G. XY(F) and YZ (G) images of border cells and junctures from a timelapse series labeled with the indicated markers. Yellow box marks the same juncture shown in F and G. White box in F shows the leading cell’s nucleus in greyscale. H. Plot showing individual juncture widths relative to time of protrusion entry (green) and nuclear entry (magenta) normalized to the size of the initial juncture. A mixed effects model (REML) followed by a tukey’s multiple comparison test was performed for statistics. N=10 egg chambers. I-J. Example images from time lapse series of border cell nuclei (I) or polar cell nuclei (J) labeled with UAS-dsRED.NLS expressed by c306Gal4(I) or UpdGaL4(J). Magenta arrowheads label the lead cell’s nucleus. K-L Plots showing individual and average +/- SEM nuclear aspect ratios (K) and changes in nuclear aspect ratio (L).An ANOVA followed by Tukey post-hoc was performed, analysis was performed on N= 10 movies for border cells and N=12 movies for polar cells. Scale bars: 10 ***μ***m. Genotypes and experimental replicates reported in Table 2.

To assess the relative sizes and dynamics of the clusters, cells, nuclei and available spaces, we acquired time series images at delamination. We labeled extracellular spaces (junctures) with fluorescent 10 kDa dextrans (Fig. 1D), nuclei with a dsRED-tagged nuclear localization signal (NLS), and border cells with a GFP-tagged actin-binding domain of moesin (Fig. 1D). Border cell clusters were ∼15-20 um in diameter, whereas individual border cells averaged ∼7.5 um, and each nucleus measured ∼5 um (Fig. 1E). The junctures that clusters initially moved into during delamination averaged 1.5 um in diameter, which was ∼3 fold narrower than even a single border cell nucleus (Fig. 1E-F). As border cells delaminated the leading protrusion widened, significantly expanding the juncture (Fig. 1F-H, S1A, Video 1). These data suggest that the leading protrusion might pry nurse cells apart before nuclear translocation.

The lead cell nucleus elongated as it moved into the protrusion (Fig. 1F, S1A) and then recovered a rounder shape as the protrusion widened (Fig. 1F, S1A-B), so nuclear shape was variable as nuclei transiently elongated into protrusions (Fig. 1F, S1B, Video 1). The width of the juncture expanded by ∼5 fold as first the protrusion and then the nucleus moved in (Fig. 1F-H). Nuclear diameter and protrusion base width tended to oscillate simultaneously when nuclei were in the protrusion (Fig S1C-D). In contrast, the tip of the spear-shaped protrusion was typically several microns narrower than the protrusion base and its size fluctuations did not correlate consistently with the base diameter (Fig S1C-E). Thus, the leading cell nucleus transiently deforms as it moves, and its translocation correlates with further expansion of the spaces between substrate nurse cells.

Lead cell nuclei tended to deform when entering the base of the protrusion (Fig. 1F and 1I, Fig. S1F, Videos 2-3), This deformation occurred in less than one minute (Fig. S1F, Video 2), unlike the hours-long process of nuclear deformation observed in breast cancer cells and fibroblasts migrating in silicone channels (Davidson et al., 2014; Denais et al., 2016). The nuclei of follower border cells also elongated and deformed though not as much as lead cell nuclei (Fig. 1K-L), and entry of follower cells further widened the path (Fig. S1G). The inner pair of nonmotile polar cells had the lowest and least variable nuclear aspect ratio (Fig. 1I-L, Video 3). These data indicate nuclear dynamics vary based on position and/or cell type, and the lead cell nucleus experiences the most nuclear deformation during collective border cell migration.

To test if transient changes in nuclear shape occurred spontaneously or due to forces resulting from migration, we measured nuclear shapes in immobile border cells expressing a dominant negative form of the small GTPase Rac (RacN17). RacN17-expressing cells lack lead protrusions and completely fail to move (Murphy and Montell, 1996). RacN17-expressing border cells had significantly lower variation in nuclear shape compared to control border cell nuclei (Fig. S1 H-K). Thus, outer border cell nuclei, particularly when leading the cluster, undergo transient shape changes as they migrate to a greater extent than inner polar cells, which may be insulated from compressive forces by the surrounding cells.

### B-type lamin promotes border cell movement between nurse cells

We next asked how nuclear lamins affect the ability of border cells to migrate into confined space. A and B-type lamins form independent intermediate filament networks that confer mechanical support to the nuclear envelope (Wintner et al., 2020; Davidson and Lammerding, 2014; Xie et al., 2016). The *Drosophila* genome encodes one A-type lamin called *LamC* and one B-type lamin named *Lam* (Bossie and Sanders, 1993; Lenz-Böhme et al., 1997) (Fig. 2A). We found that both Lam (Fig. 2B) and LamC (Fig. 2C) antibodies stained the periphery of border cell nuclei, and staining was diminished in cells expressing the corresponding RNAi (Fig. 2D-G and S2A-G). Expressing two different UAS-RNAi lines, referred to as Lam1 RNAi and Lam2 RNAi, which target separate regions of the Lam gene, with a Gal4 that drives expression in border and polar cells (c306Gal4), caused migration defects (Fig. 2D-E, and 2H-I). In contrast, LamC RNAi did not (Fig. 2F-G and 2J). Gal4 is more active at 29 °C than at lower temperatures. When we incubated Lam RNAi-expressing flies at 29 ºC for 1 day, we observed variable knockdown, and the clusters with the most effective knockdown exhibited more severe migration defects (Fig. S2H). When we incubated flies for 3 days at 29º C, the lamin depletion was stronger (Fig. S2F). Migration defects were also stronger: 82% of clusters expressing Lam RNAi line Lam1RNAi, and 72% of clusters expressing Lam RNAi line Lam2RNAi showed incomplete migration at stage 10. In both lines, half of border cell clusters failed to move in between nurse cells (Fig. 2I).

**Figure 2.**
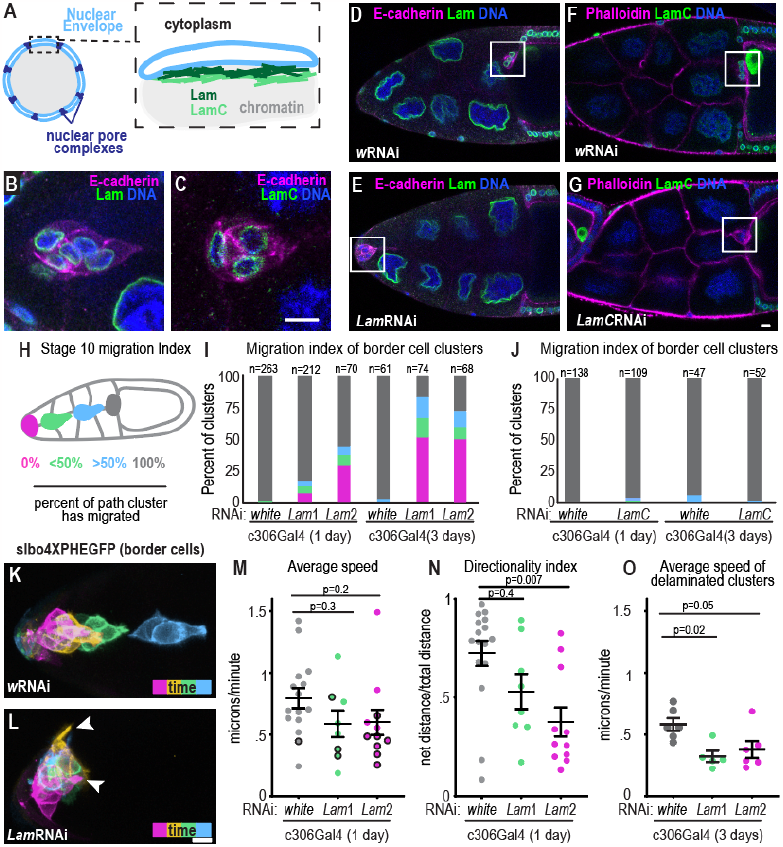
B-type lamin promotes border cell delamination. A. Schematic of the nuclear envelope and the two *Drosophila* lamin proteins, Lam and LamC. B-C. Images of fixed border cell clusters stained with Lam (B) and LamC (C). Scale bar:10 microns. F. Schematic of migration index parameters. D-G. Images of fixed stage 10 egg chambers for the indicated conditions images shown are 1 day at 29 ºC incubation. The oocyte is always shown on the right and the white box marks the border cell cluster. H. Schematic of migration index used to score stage 10 egg chambers. I-J Plots showing migration indexes for indicated conditions. n=number of egg chambers. I. Statistical test, 1 day at 29 ºC: Fisher’s exact test with Bonferroni correction yield significant difference in % with complete migration, wRNAi vs Lam1RNAi, p<0.0002, and wRNAi vs Lam2RNAi, p<0.0002. Statistical test, 3 day at 29 ºC: Fisher’s exact test with Bonferroni correction for difference in % with complete migration, wRNAi vs Lam1RNAi, p<0.0002, and wRNAi vs Lam2RNAi, p<0.0002. J. Statistical test 1 day at 29 ºC: Fisher’s exact test for difference in % with complete migration in wRNAi versus LamCRNAi 1 Day, p=0.08, and Fisher’s exact test wRNAi versus LamCRNAi 3 day yields p=0.3. K-L Maximum projections of time frames from time lapse series of stage 9 egg chambers expressing slbo4xPHEGFP to mark border cell membranes in control (wRNAi, K) and Lam-depleted border cells (LamRNAi, L) after 1 day at 29 ºC. Time points relative to the start of imaging: magenta=0 minutes, yellow=33 minutes, green=54 minutes, blue=108 minutes. Arrowheads mark protrusions. M. Cluster speed of individual (dots) and mean +/- SEM (bars) control and Lam-depleted clusters. Black outlines: border cell clusters that did not move away from anterior during imaging session. Kruskal-Wallis Test. N. Individual and mean +/- SEM directionality index of clusters measured as total path traveled/net distance. Kruskal-Wallis Test. O. Individual and mean +/- SEM speed of delaminated clusters after 3 day incubation at 29 C. An ANOVA followed by Tukey post-hoc was performed for statistical testing. Genotypes and experimental replicates reported in Table 2.

To assess how lamins affect border cell motility and behavior, we performed live imaging of stage 9 egg chambers. During delamination, lead border cells from controls rounded up and extended one main protrusion in between nurse cells before moving in between the nurse cells towards the oocyte (Fig. 2K, Video 4). In contrast, Lam-depleted clusters extended short-lived and ectopic protrusions (Fig. 2L and Video 4). Clusters were mobile and sometimes moved between anterior follicle cells and germ cells instead of taking their normal path between nurse cells (Fig. 2K-M). However Lam-depleted clusters exhibited less directional persistence (Fig. 2N). In the 1-day RNAi-treated clusters, movement away from the anterior end of the egg chamber was significantly delayed (Fig. S2I). Clusters that delaminated exhibited similar migration speeds to controls (Fig. 2M), likely due to incomplete knockdown though it was not possible to assess knockdown efficiency in living samples. With a 3-day incubation, the delamination defect was more penetrant and those lamin-depleted clusters that did delaminate migrated slower (Fig. 2O). Together we conclude that a partial Lam knockdown delays delamination, whereas a more severe Lam-depletion hinders delamination and slows migration.

To test if Lam is required in the outer border cells or polar cells, we crossed UAS-Lam RNAi lines to fruitlessGal4, which is expressed in border but not polar cells, and UpdGal4, which is expressed in polar but not border cells. LamRNAi expressed with fruitlessGal4 but not with UpdGal4 caused incomplete migration at stage 10 (Fig. S2 J-K). We conclude the *Drosophila* B-type lamin is required in the outer, motile border cells.

### B-type lamins are required to maintain nuclear shape and integrity

Depletion of either A-type or B-type lamin disrupts the nuclear envelope permeability barrier in cultured cells (Vargas et al., 2012). To test how Lam and LamC depletion affect border cell nuclei, we performed live imaging of dsRED.NLS, which is retained in the nucleus when the nuclear permeability barrier is intact. Control (Fig. 3A, A’) and LamC-depleted (Fig. 3B, B’) nuclei, dsRED remained inside the nucleus in all cells observed. By contrast, some Lam-depleted cells exhibited dsRED.NLS throughout the cell (Fig. 3C-D). Live imaging also revealed that a subset of Lam-depleted leading cell nuclei in protrusions herniated backwards, whereas neither LamC-depleted nor control nuclei exhibited such herniations (Fig. 3E-G). Lam-depleted nuclei within protrusions were even more elongated than controls (Fig. 3H), consistent with the proposed role of B-type lamins in nuclear elasticity (Wintner et al., 2020; Harada et al., 2014). A subset of leaders had reduced circularity in Lam-depleted but not LamC-depleted nuclei (Fig. 3I). Lam-depleted nuclei were also slightly smaller than controls (Fig. 3 J-L) though cell area was not affected (Fig. 3M). We conclude that Lam is required to maintain border cell nuclear integrity, shape, and size, while LamC is less critical.

**Figure 3.**
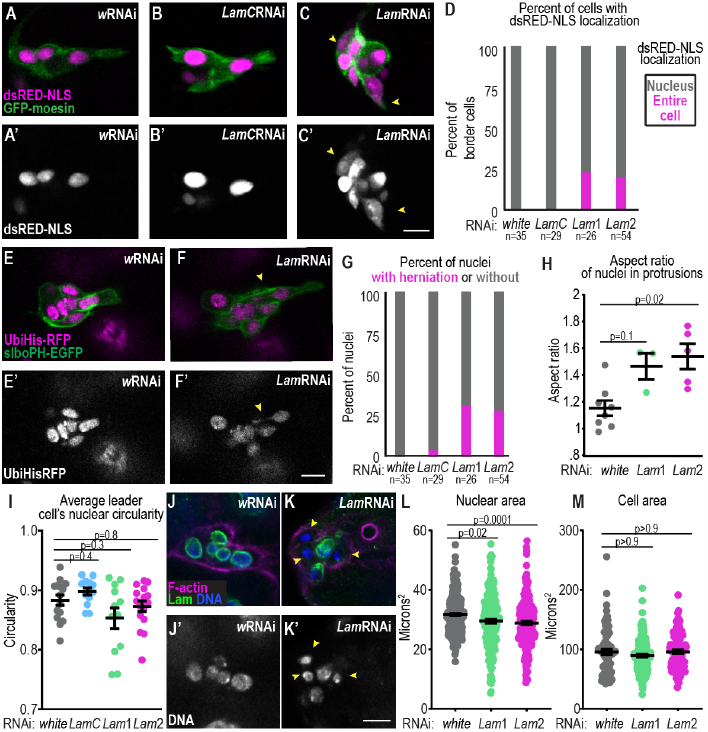
B-type lamins are required to maintain nuclear shape and integrity. A-C. Images of border cells from time lapse series in stage 9 egg chambers with the indicated markers and conditions. Yellow arrowheads point to cells with dsRED.NLS throughout the cell. D. Plot measuring the percentage of cells with dsRED.NLS throughout the cell versus in the nucleus. A Fisher’s exact test with a Bonferroni correction for multiple testing was performed: (w vs. Lam1 p=0.01), (w vs Lam2,p=0.009) (w vs. LamC, p>0.9). n=number of nuclei. E-F. Images of border cells from time lapse series with the indicated markers and conditions. Yellow arrowhead marks backward nuclear herniation. G. Plot with the percent of nuclei with herniations in each condition. A Fisher’s exact test with a Bonferroni correction for multiple testing was performed: (w vs. Lam1 p=0.001), (w vs Lam2,p=0.0009) (w vs. LamC, p>0.9). n=number of nuclei. H. Plot of individual and mean +/- SEM nuclear aspect ratios when the nucleus extended into the protrusion (defined as >1 SD above average nucleus distance to polar cell boundary). A one way ANOVA with post-hoc Tukey was performed I. Plot of individual and average +/- SEM values for nuclear circularity. A Kruskal-Wallis test was performed. Dots show # of leader cell nuclei analyzed. J-K Images of fixed border cell clusters stained with indicated markers for wRNAi (J) and LamRNAi (K) border cells. L-M Measurements of nuclear area and cell area. Each individual area value shown as dots and bars representing mean +/- SEM. Statistical test L-M: Kruskal wallis test. Scale bars: 10 ***μ***m. Figure 3 KD condition: 1 day at 29 ºC. except the cell area (M) which was 3 days at 29 ºC. Genotypes and experimental replicates reported in Table 2.

### Effects of Lam and LamC overexpression on border cell migration

In some contexts, A-type lamins impede migration (Davidson et al., 2014; Rowat et al., 2013), however, overexpression of LamC did not disrupt migration (Fig. 4A-B). While overexpression Lam with a RNAi-resistant version of Lam (LamRE) also did not impede migration, stronger overexpression of Lam with a different construct caused migration defects (Fig. 4A-F). Strong Lam overexpression perturbed nuclear shape (Fig. 4D), consistent with the proposed role of B-type lamins in regulating nuclear membrane abundance (Prüfert et al., 2004). Expression of the re-encoded, RNAi-resistant Lam construct (LamRE) fully rescued Lam RNAi migration defects (Fig. 4G-H). LamC overexpression partially rescued Lam RNAi migration defects (Fig. 4I-K). We conclude that optimal Lam levels are critical for border cell migration.

**Figure 4.**
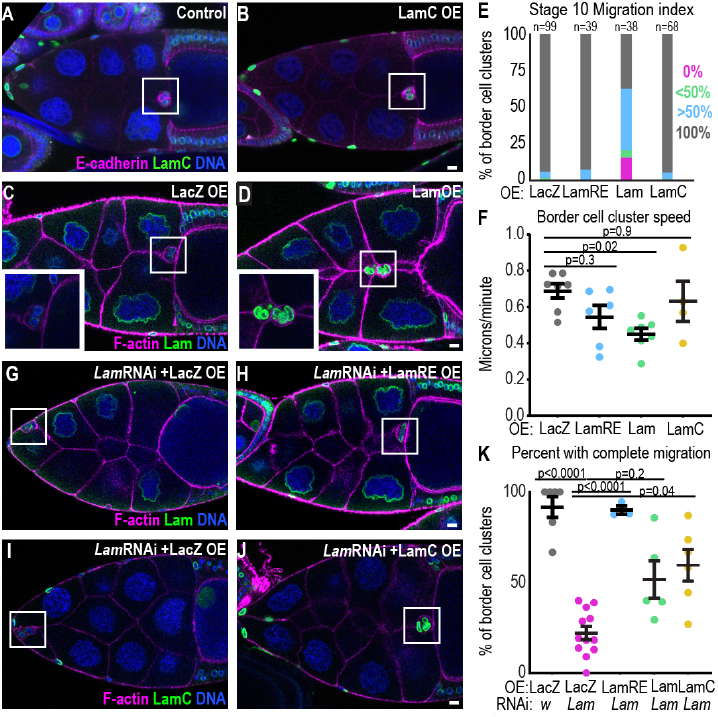
Lamin overexpression compensates for LamRNAi but strong Lam overexpression impedes border cell migration. A-D. Representative images of stage 10 egg chambers for the indicated conditions. White box marks border cell clusters. E. Migration indices for the indicated conditions. n=number of egg chambers. A Fisher’s exact test with a Bonferroni correction to test for proportion with complete migration yields a significant result between LacZ versus Lam OE, p< 0.0003. All other comparisons to LacZ are not significant. F. Plot of cluster speed with individual movies and mean +/- SEM plotted. Statistical test: One way ANOVA followed by post-hoc Tukey. G-J. Representative images of stage 10 egg chambers for the indicated conditions. White box surrounds border cell cluster K. Plot of the mean +/- SEM percentages of egg chambers with completed migration, each dot represents one experimental replicate (N). n (number of egg chambers): wRNAi/LacZ: 47 LacZ/LamRNAi: 290 LamRNAi/LamRE: 59 LamRNAi/LamOE: 73 LamC/LamRNAi: 149. Statistical test: Brown-Forsythe and Welch followed by Dunnett’s Multiple Comparison test. Genotypes and experimental replicates reported in Table 2. Scale bars: 10 ***μ***m.

To further test the effects of perturbing nuclear structure on border cell migration, we investigated the effects of overexpressing the Lamin B receptor (LBR). LBR is an inner nuclear membrane protein that is responsible for the multilobed nuclei that are proposed to allow neutrophils to squeeze into and out of blood vessels and tissues (Hoffmann et al., 2002). LBR overexpression causes irregular nuclear morphology in *Drosophila* embryos (Hampoelz et al., 2016) and interferes with the nuclear ruler mechanism in HeLa cells (Lomakin et al., 2020). In control (Fig. S2L) and LBR-overexpressing border cells (Fig. S2M), LBR localized at the nuclear periphery. Nuclei overexpressing LBR became elongated (Fig. S2N-P). To test if LBR overexpression disrupted migration in border cells or polar cells, we crossed UAS-LBR and a control (UAS-LacZ) to fruitlessGal4 and UpdGal4. Overexpression of LBR with fruitlessGal4 but not UpdGal4 caused significant migration defects, indicating LBR OE affects the outer, motile border cells (Fig. S2Q-R). Thus, LBR overexpression disrupts nuclear shape and impedes border cell movement between nurse cells, supporting that border cells require specific nuclear properties and biochemical composition to move within their naturally confined environment.

### B-type lamin promotes expansion of the lead protrusion

To assess how Lam depletion might affect border cell invasion between nurse cells, we first determined if LamRNAi-expressing border cells are specified normally. Border cell fate specification, cluster formation, and movement require the activity of multiple transcription factors and their downstream targets (Montell et al., 2012). In addition to their mechanical roles, lamins are required for chromatin organization, gene expression, and cell survival (Davidson and Lammerding, 2014; Chen et al., 2019; Harada et al., 2014). So, we tested the effect of Lam RNAi on border cell fate, cluster formation, and expression of genes required for initiation of migration. Lam-depleted border cells still formed clusters (Fig. S3A-C) with similar circularity (Fig. S3D) and cell numbers (Fig. S3E) as controls. Further, Lam-depleted clusters had similar F-actin (Fig. S3F) and E-cadherin (Fig. S3G-H) levels to controls. STAT activation, which is essential for border cell specification (Silver and Montell, 2001), was actually increased ∼2-fold in Lam-depleted cells at stage 8 (Fig. S3I-K), which is consistent with reports that Lam and the nuclear-membrane-associated protein, Dysfusion, limits STAT signaling (Wu et al., 2022; Petrovsky and Großhans, 2018). Increased STAT in border cells does not impede their migration (Silver and Montell, 2001; Silver et al., 2005). Ectopic STAT can cause additional border cell clusters to form, however we did not observe extra border cells in Lam depleted clusters suggesting that the elevated STAT signaling was insufficient to induce extra border cells. Notch activity, which is required for border cell delamination (Wang et al., 2007), was similar to controls (Fig. S3 L-M). We conclude that lamin is not required for border cell specification.

Next we investigated how lamins affect cluster polarity given some reports that the nucleus is required for cell polarization while others show that polarity can develop even in enucleated cells (Graham et al., 2018; Lee et al., 2007). Border cells maintain three types of polarity that are important for migration: 1) Apical-basolateral polarity 2) inside outside polarity and 3) front-back polarity (Montell et al., 2012; Pinheiro and Montell, 2004; Wang et al., 2018; Duchek et al., 2001; McDonald et al., 2008; Assaker et al., 2010; Luo et al., 2019). Polar cells had an apical enrichment of E-cadherin and were found on the inside of control (Fig. S4A) and Lam-depleted clusters (Fig. S4B). E-cadherin was also enriched at cell-cell junctions in both conditions (Fig. S3A-C, and S4A-B). Similar to controls (Fig. S4A and C), Lam RNAi clusters (Fig. S4B and D) had lateral localization of Discs large (Dlg) and apical enrichment of Atypical Protein Kinase C (aPKC). Further, control (Fig. S4E) and LamRNAi clusters (Fig. S4F) displayed enrichment of F-actin on the outside of the cluster compared to inside and in forward directed protrusions (Fig. S4 G-I), although Lam-depleted clusters also formed ectopic side protrusions (Fig. 2L). These data indicate border cells retain features of cluster polarity upon Lam-depletion.

The observations that nuclei normally move into protrusions, which then widen, and that Lam depletion impedes delamination and migration without affecting cluster specification or polarity suggested that border cell nuclei, in particular the lead cell nucleus, might function as a wedge to stabilize and enable enlargement of the lead protrusion. To explore this possibility further, we compared nuclear shape changes to protrusion shape changes over time. In controls, the movement of leading cell nuclei into protrusions corresponded with a widening of the protrusion neck (Fig. 5A, Video 5). In lamin-depleted clusters, nuclei extended forward, but did not remain in the protrusion neck, and protrusions narrowed (Fig. 5B, Video 5). On average, protrusions were shorter and thinner in LamRNAi-expressing cells compared to controls (Fig. 5C-D). In control leading cells, nuclear movement forward into the protrusion was associated with protrusion widening, while backward movement correlated with protrusion narrowing (Fig. 5E). In Lam-depleted cells, protrusion width was not significantly changed by nuclear movement (Fig. 5E). Further, nuclear width and protrusion base width did not correlate as well upon Lam depletion compared to control clusters (Figure 5F-J). We conclude that Lam is required to maintain and facilitate expansion of the leading protrusion.

**Figure 5.**
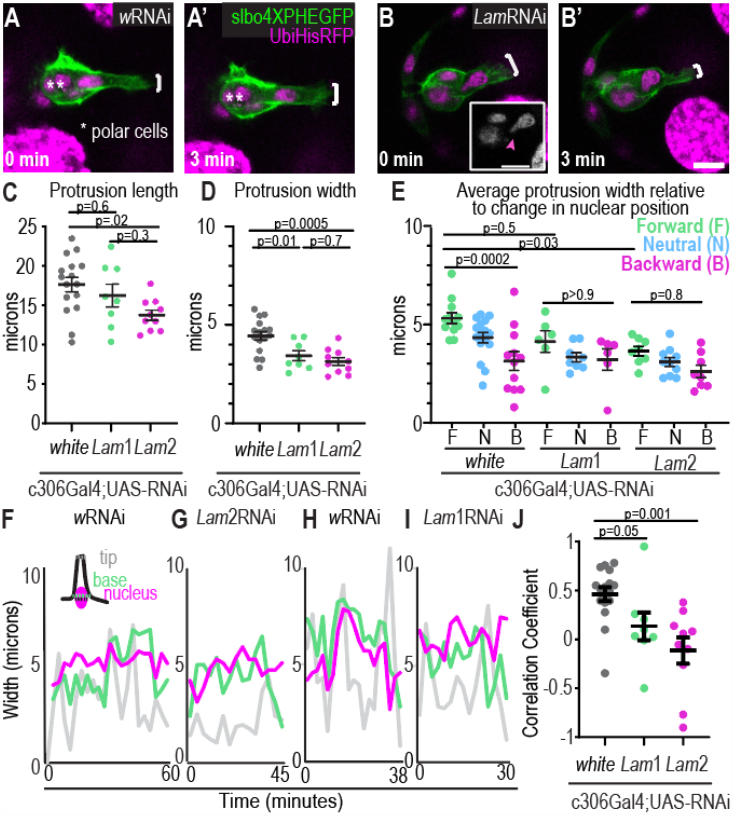
B-type lamins promote the expansion of a single protrusion. A-B Single z-slice images from time lapse series of migrating control border cells (wRNAi, A) or Lam-depleted border cells that delaminate but fail to expand the protrusion (B). White bars show protrusion width. C-E Plots showing average +/- SEM (bars) and individual movie values (dots) of the average length (C) and width of protrusions (D-E) for the indicated conditions. G. shows average protrusion width relative to nuclear movement for each RNAi line. An ANOVA with Tukey post-hoc was performed for each plot. F-I Example plots showing the width of the protrusion tip, base, and nucleus for the indicated conditions for an individual cluster over time. J. Plot with dots of individual and mean +/- SEM correlations between nuclear width and base width for each cluster’s first protrusion. A Kruskal-Wallis Test was performed. N= 16 (wRNAi), N=8 (Lam1RNAi), and N=10 (Lam2RNAi) movies for each condition shown in C-E. and J. Scale bars: 10 ***μ***m. All conditions are after 1 day incubation at 29 ºC. Genotypes and experimental replicates reported in Table 2.

The result that Lam promotes protrusion expansion is, in principle, consistent with the nuclear piston mechanism (Lee et al., 2021). One feature of the piston effect is that there is a pressure gradient and diffusion barrier between the front and back of the confined cell (Petrie et al., 2014). So we tested for a diffusion barrier in the leading cell by illuminating a photoactivatable GFP-*α*tubulin in at the front of a leading border cell protrusion in front of the nucleus and then measuring its diffusion to the opposite side of the nucleus at the back of that cell (Fig. 6A, Video 6). GFP diffused rapidly (Fig. 6A). For comparison, we photoactivated at the basal side of epithelial follicle cells, which should not have a diffusion barrier, and assessed diffusion to the apical side of the nucleus (Fig. 6B, Video 6). The ratio of GFP fluorescence between activated:unactivated regions were similar between the follicle cell and border cell over time (Fig. 6C), and diffusion time constants in unactivated regions were not significantly different between protruding border cells and epithelial follicle cells (Fig. 6D). We conclude that border cells lack a key hallmark of the nuclear piston.

**Figure 6.**
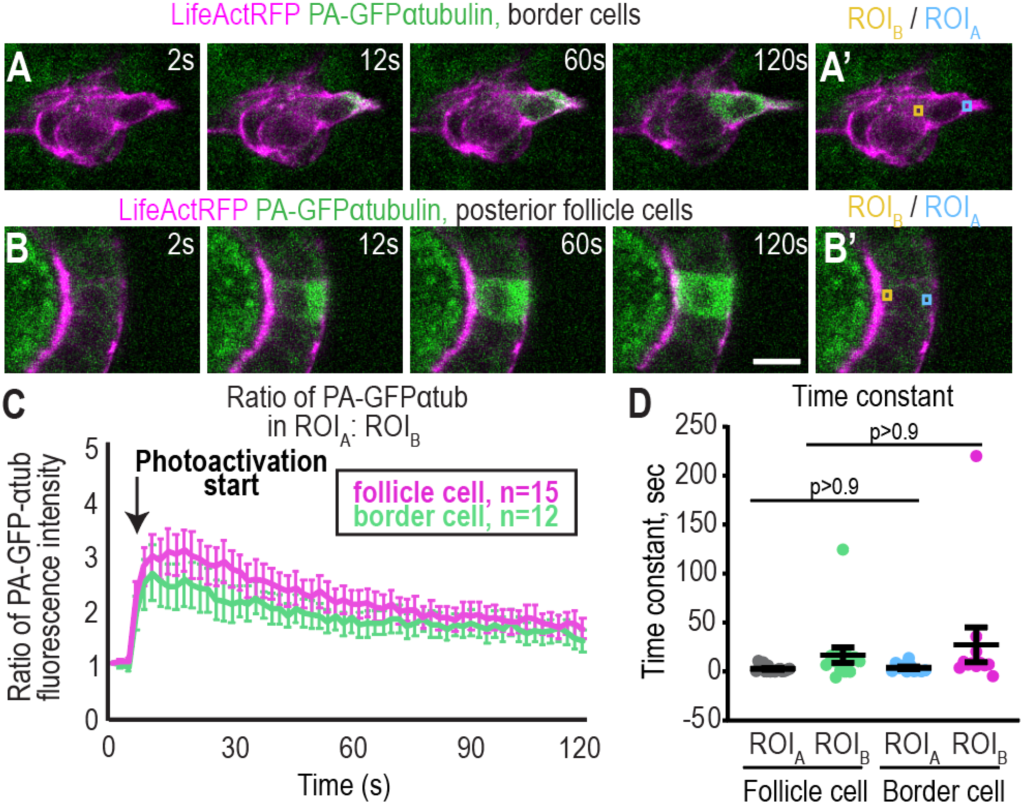
There is no detectable diffusion barrier in the leading border cell. A-B Images from time lapses series where GFP-αtubulin was activated in a border cell (A) or a posterior follicle cell (B). A’-B’ show where regions of interest (ROIs) were measured. Scale bars: 10 ***μ***m. Time is relative to the start of the movie and photoactivation occurred at 6 seconds. C. Plot showing the mean +/- SEM of the ratio of the fluorescence intensity of GFP in front:back ROIs over time. Experiments were performed on 3 different experimental days and n=number of movies is displayed for each condition. D. Time constants calculated from the slope of Boltzmann sigmoidal fitted curve (see methods). A Kruskal Wallis test was performed. Genotypes and experimental replicates reported in Table 2.

### Myosin II cortical flashes correspond with nuclear movement and shape changes

An additional mechanism by which lamins may promote border cell delamination is through the nuclear ruler. The nuclear ruler model proposes that when cells are confined to a space narrower than the nuclear diameter, the nuclear envelope stretches out, leading to calcium release and recruitment of cytosolic phospholipase A2 (cPLA2) to the nuclear membrane. This calcium initiates a signaling cascade that recruits myosin to the cell cortex in many cell types including HeLa and Zebrafish cells (Lomakin et al., 2020; Venturini et al., 2020), and stimulates massive blebbing, which results in cells escaping the confinement. Border cells do not normally exhibit blebs, and there is no identifiable cPLA2 encoded in the fly genome (Ben-David et al., 2015). There is a calcium-independent iPLA2, so we tested its effect on migration. Neither of two independent null mutants (Lin et al., 2018) exhibited any border cell migration defect (Fig. S5A). We also used GCaMP to evaluate calcium dynamics in border cells. While earlier stage follicle cells show rapid pulses of calcium with the GCaMP sensor as reported previously (Sahu et al., 2017), migrating border cells did not exhibit spatial or temporal changes in GCaMP fluorescence (Fig. S5B-E, Video 7). Therefore, border cells lack key features of the nuclear ruler mechanism.

We then tested the effects of lamins on myosin II dynamics because myosin II is essential for border cell migration. Myosin II accumulates at the cortex of border cell clusters in transient foci or flashes (Majumder et al., 2012). Myosin II flashes at the back help clusters delaminate (Majumder et al., 2012) while flashes on the side and front retract protrusions (Mishra et al., 2019b) and counteract nurse cell compression (Aranjuez et al., 2016). We used a fluorescently tagged myosin light chain (Sqh-mCherry) to observe the effect of Lam RNAi on myosin. In both control (Fig. 7A, Fig. S5F) and Lam-depleted (Fig. 7B, Fig. S5G) clusters, Sqh-mCherry appears transiently in “cortical flashes.” Myosin flashes occurred at similar frequencies in control and Lam-depleted clusters and throughout the cortex (Fig. 7A-C, Fig. S5 F-G, Video 8), indicating that Lam is not required for myosin recruitment to the cortex.

**Figure 7.**
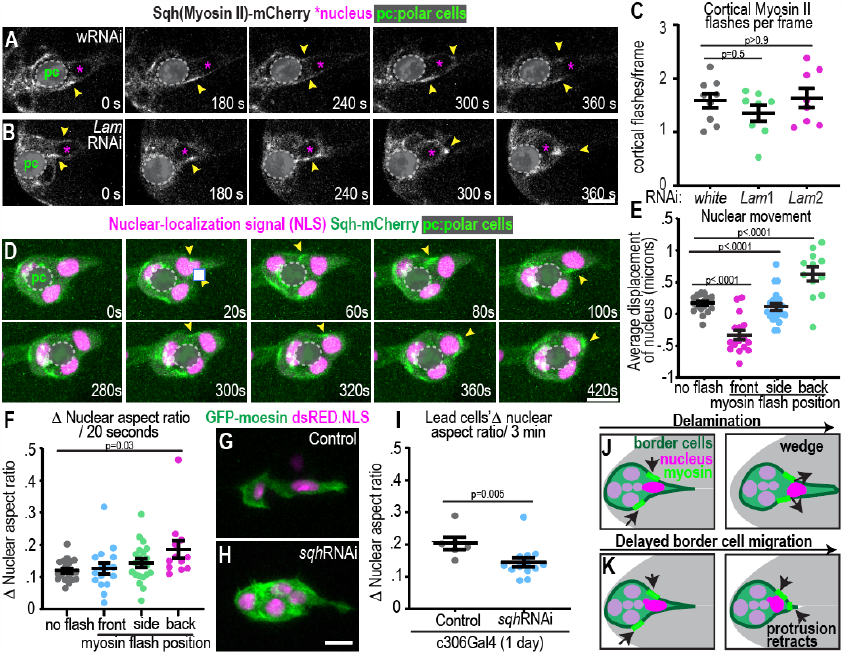
Myosin II cortical flashes correspond with nuclear movement and shape changes. A-B Example images of Sqh-mCherry in control (A) and Lam-depleted clusters (B) after 1 at 29 ºC (see also Fig. S5F-G for 3 day 29 ºC). Sqh-mCherry aggregates at apical surfaces in polar cells(pc) as previously reported (Mishra et al., 2019b) and has been covered to focus on cortical flashes. White arrowheads point to Sqh flashes. C. Average number of myosin flashes observed divided by the total number of frames images RNAi condition: 3 day 29 ºC. An ANOVA with post-hoc Tukey was performed. D. Images from a timelapse series showing myosin flashes around the nucleus. Yellow arrowheads point to flashes. E-F. Average change in nuclear position along the A-P axis of the egg chamber (E), and average nuclear aspect ratio change (F) relative to presence and position of myosin flashes. N=16 movies, each dot represents one nucleus, bars: mean+/-SEM. Statistical tests E-F: One way ANOVA followed by Tukey post-hoc. G-H. Images of control and Sqh-depleted clusters showing that nuclei stay behind protrusion necks upon SqhRNAi, 1 at 29 ºC. I. Plot of the change in nuclear aspect ratio of leader cells. Dots: individual nuclei, middle bar and error bars: mean +/- SEM. 1 at 29 ºC. Each dot represents one nucleus. Statistical test: Mann-Whitney Test. J-K Working model for how nuclei and myosin coordinate delamination and invasion into confined space. Genotypes and experimental replicates reported in Table 2. Scale bars: 10 ***μ***m.

In control clusters, cortical myosin flashes correlated with nuclear movement. When myosin accumulated behind the nucleus it moved forward into the protrusion, and the nucleus changed shape; in contrast, flashes in front of the nucleus correlated with backward nuclear movement (Fig. 7D-F, Videos 2 and 9). Myosin flashes accumulated behind the lead cell nucleus in both control and Lam-depleted clusters (Fig. 7A-C, Fig S5F-G, Videos 8-9). In lamin-depleted clusters with failed delamination, rather than pushing the nucleus forward, the flashes moved ahead of the nucleus and the protrusion retracted (Fig. 7B, S5G).

To test the functional significance of myosin on nuclear movement, we expressed Sqh RNAi and evaluated the effect on nuclear shape and movement. A partial Sqh-depletion reduced leader cell nuclear shape changes during migration (Fig. 7 G-I). A more penetrant Sqh depletion resulted in long and long-lived protrusions and failed delamination (Video 10) as reported previously (Majumder et al., 2012; Mishra et al., 2019b). In these clusters nuclei failed to enter the protrusion (Fig. 7H and Video 10). We conclude myosin forces deform and position nuclei.

Nuclei are connected to the cytoskeleton through the Linker of the Nucleoskeleton and Cytoskeleton (LINC) complex (Crisp et al., 2006). However, prior work suggests nuclei in glioma cells can be deformed and moved directly by myosin, specifically when moving in 3D through pores smaller than the nuclear diameter (Jahed and Mofrad, 2019; Lee et al., 2021). So we tested if LINC complex components were involved in border cell migration. Depletion of LINC complex components, Klar, Koi, or Msp300 with previously tested RNAi lines (Collins et al., 2017; Perillo and Folker, 2018), did not cause delamination or major migration defects (Fig. S5H-J). These data suggest myosin flashes could be directly acting on nuclei. Together the data suggest that cortical myosin flashes may push nuclei into protrusions, and Lam is required for the protrusion to grow appropriately and facilitate movement into the tiny spaces between nurse cells (Fig. 7J).

## Discussion

*In vivo*, cells migrate through diverse terrains: they can move on basement membranes or through ECM gels; but they can also squeeze between cells or in spaces between ECM and cells (Mishra et al., 2019a; Rørth, 2009; Yamada and Sixt, 2019). An important difference between cell movement through matrix versus between cells is that migrating cells can secrete proteases to enlarge spaces in matrix (Wolf et al., 2013; Yamada and Sixt, 2019; Wolf et al., 2007). However, cells migrating between other cells, for example during transendothelial migration or during embryonic development, do not have the option to degrade the substrate. Instead, they must either deform themselves sufficiently to squeeze through or pry the substrate cells apart. How cells and especially cell collectives achieve this is not well understood. Here, using the border cells as a model, we report the dynamics of nuclei and discover a novel role for the nucleus in this collective, confined, *in vivo* cell migration.

We propose a nuclear wedge model (Fig. 7J), whereby F-actin-rich protrusions begin to separate nurse cells by breaking nurse cell-nurse cell adhesions. Then, the nucleus moves into the protrusion, preventing its collapse and allowing it to enlarge. Myosin flashes at the cortex appear to push the lead cell nucleus into the protrusion during successful delamination. However, when these flashes move past the nucleus, they retract the protrusion, impeding forward invasion (Fig. 7K).

The leading border cell shares features of confined single cells migrating *in vitro* and *in vivo*. In cells migrating in confined hydrogels, in neuronal progenitor cell-on-cell migrations, and in glioma cells invading the neural cortex, nuclei are pushed from behind by myosin similar to what we observe in the leading border cell (Norden et al., 2009; Lee et al., 2021; Beadle et al., 2008). Also like border cells, there does not appear to be a role for canonical components of the LINC complex in these movements. In confining hydrogels and our system, nuclei facilitate protrusion expansion. We sometimes observe transient expansion of the protrusion ahead of the nucleus reminiscent of a piston effect, and we cannot directly measure pressure in front of the nucleus *in vivo*, so it is not possible to rule out the possibility that the nucleus may function as a piston. However other nuclear piston hallmarks (Petrie et al., 2014), such as the diffusion barrier, requirement for the LINC complex and connection to intermediate filaments are missing from border cells. The piston effect also requires myosin activity in front of the nucleus for protrusion expansion, whereas in border cells, myosin activity behind the nucleus correlates with protrusion expansion. Protrusion expansion ahead of the nucleus in border cells might instead be due to other factors including F-actin polymerization since Rac activity and F-actin are elevated in protrusions (Murphy and Montell, 1996).

Hallmarks of the nuclear ruler were also absent from border cells including any requirement for PLA2, calcium fluctuations, or blebbing. The experiments that uncovered the nuclear ruler severely compressed cells using cantilevers that were themselves not deformable, whereas border cells push nurse cells aside. This difference in the source of compression and material properties of the microenvironment may contribute to differences in cellular responses. The relatively mild deformations of lead border cell nuclei at least superficially most resemble those of proteolytically active cells migrating through dense matrix, which can relieve compressive forces by matrix degradation. Perhaps the ability to deform nurse cells similarly reduces the mechanical load on border cells.

Lamins enable expansion of the leading border cell protrusion and movement into confined space. Lam-depleted nuclei become more elongated when squeezed within protrusions and herniate backwards (Fig. 3), indicating that Lam counteracts rearward forces, likely generated by myosin-mediated contractions and/or nurse cell compression. Lamin may have additional roles in border cells, such as effects on transcription, although we did not detect effects on expression of key border cell genes sufficient to explain the delamination defects.

Lead cell protrusions serve multiple functions in migrating border cells. They are sensory structures that probe for chemoattractants and available space (Dai et al., 2020). The data presented here suggest an additional function for the lead cell protrusion – physical enlargement of the available space in front of the cluster. The idea that the protrusion can physically expand space has been observed in hydrogels *in vitro* (Lee *et al*., 2021) and is supported by our *in vivo* study. The thin, short and unstable protrusions observed in Lam-depleted cells suggest that the mechanical and/or biochemical properties of nucleus at the base of the protrusion prevent its collapse and thereby facilitate protrusion enlargement, which is essential for the cluster to advance into the tiny spaces between nurse cells.

Depletion of Lam results in border cells with thin and ectopic protrusions (Fig. 2, 5), indicating that the B-type lamin at the nuclear envelope promotes enlargement and stabilization of the lead cell protrusion. Previous work shows that mechanical feedback from the dominant lead protrusion inhibits side and rear protrusion (Cai et al., 2014; Mishra et al., 2019b). Similar to an E-cadherin depletion, some lamin-depleted clusters have a reduced directionality index and migrate towards the side of the cluster. Without maintenance of a single protrusion towards the center of the egg chamber in Lam-depleted cells, side protrusions extend, and clusters either fail to migrate or attempt to take an abnormal trajectory (Fig. 2).

Some cells, like border cells, use the nucleus to counteract environmental confinement, while other cells, such as neutrophils, reduce lamin expression instead to squeeze through tight spaces. Still other cells overcome the barrier of confinement, not by modulating the properties of the nucleus, but rather those of the cortex. For example, the transcription factor, Dfos, controls cortical actin levels to counteract tissue compression and permit nucleus movement during macrophage invasion into spaces between the ectoderm and mesoderm in Drosophila embryos (Belyaeva et al., 2022). In these cells, lamin levels do not normally affect migration, but reduced lamin can rescue invasion when cortical tension is impaired, illustrating how cells can use multiple mechanisms to move through confined environments. Whether lamins promote migration in confinement, impede it, or are irrelevant seems to depend on a combination of the mechanical properties of the nuclei and cortex of the migrating cells as well as the microenvironment. For example, egg chambers are estimated to be 100 fold softer than the silicone (PDMS) used to fabricate microchannels (Johnston et al., 2014; Lamb et al., 2021). It may be that nuclear stiffness would be insufficient to overcome such a stiff barrier but sufficient to function as a wedge in softer environments.

We found a role for B-type but not A-type lamin in border cell migration. A- and B-type lamins contribute to nuclear stiffness (Wintner et al., 2020) and have been reported to affect cell migration in prior studies (McGregor et al., 2016). However, we find that the sole A-type lamin in Drosophila, LamC, is not essential in border cells. It is possible that this is because LamC is normally expressed at a lower level than Lam, which would explain how overexpression of LamC can partially rescue Lam RNAi even though LamC RNAi does not cause a phenotype. Expression levels of A-type lamins tend to scale with tissue stiffness and are important in stiff environments, while B-type lamins dominate in softer environments (Swift et al., 2013). The egg chamber is a relatively soft environment, consistent with the pattern that B-type lamins might be more critical, similar to its role in neural tissues, where A-type lamins are downregulated (Jung et al., 2012). An alternative, but not mutually exclusive, hypothesis is that B-type lamin might have a unique function compared to A-type lamin. This might explain why A-type lamin can only partially compensate for the loss of the B-type lamin.

Follower border cells undergo less severe nuclear deformations compared to leader cells. Follower cells contribute to overall cluster motility (Campanale et al., 2022) and further expand the opening between nurse cells as the cluster advances. Polar cells undergo very little deformation. Since severe deformations can result in nuclear rupture and DNA damage (Irianto et al., 2017; Denais et al., 2016; Raab et al., 2016), absorption of the brunt of nuclear deformation by the lead cell could in principle serve to protect the rest of the cells in a moving collective and thereby provide a selective advantage to collective movement, which might be particularly relevant in the context of tumor metastasis.

## Materials and Methods

### Drosophila genetics

*Drosophila* stocks used in this study are described in Table 1. Flies were maintained in cornmeal-yeast food. To combine Gal4s with different UAS-lines, 5-7 virgin females were crossed to 2-3 males in the presence or absence of the temperature-sensitive repressor Gal80^TS^; all progeny used were heterozygous for constructs. For lines without Gal80^TS^, crosses were maintained at 25 °C and flies were transferred into new vials every 2-3 days.

**Table 1.**
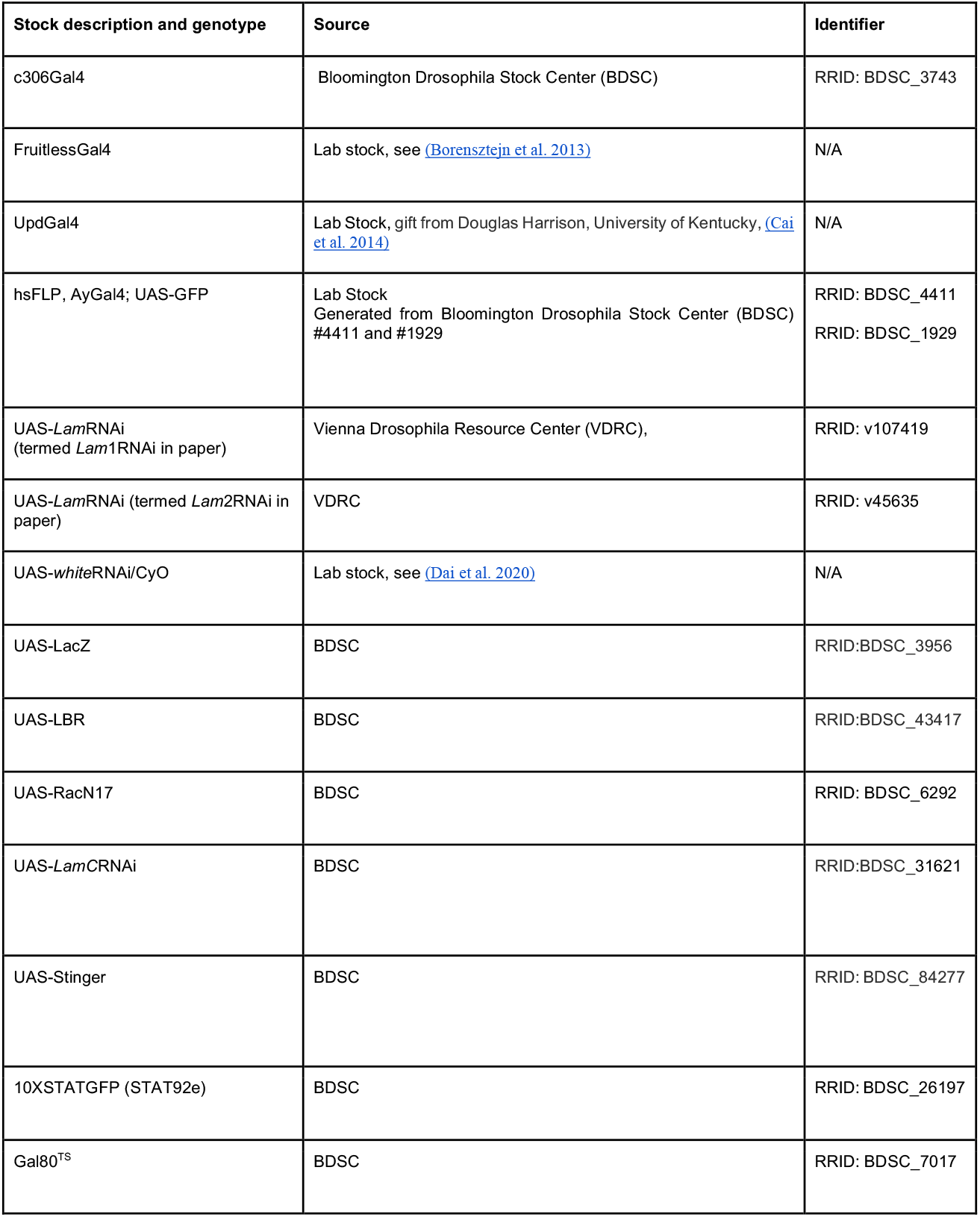
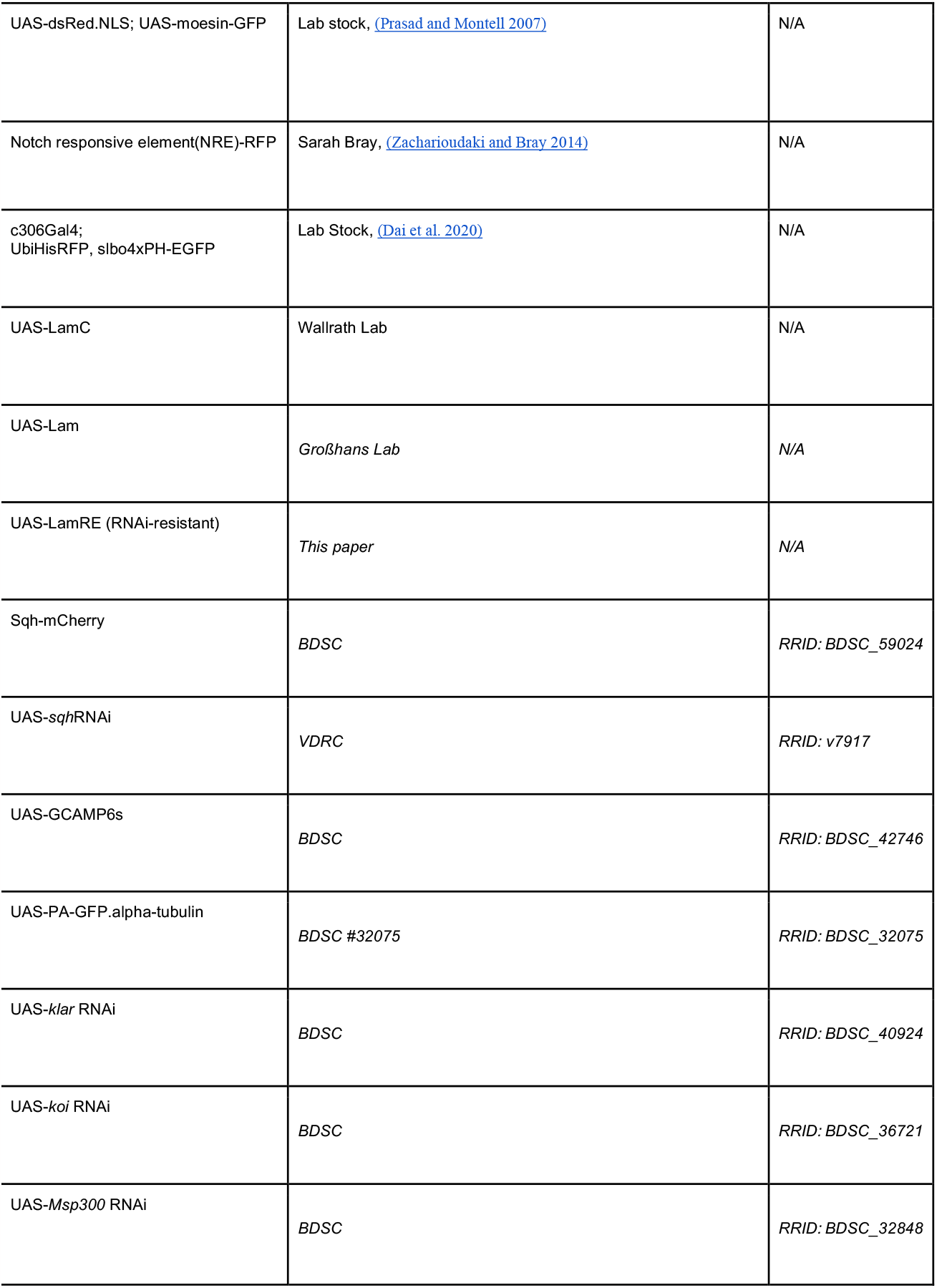
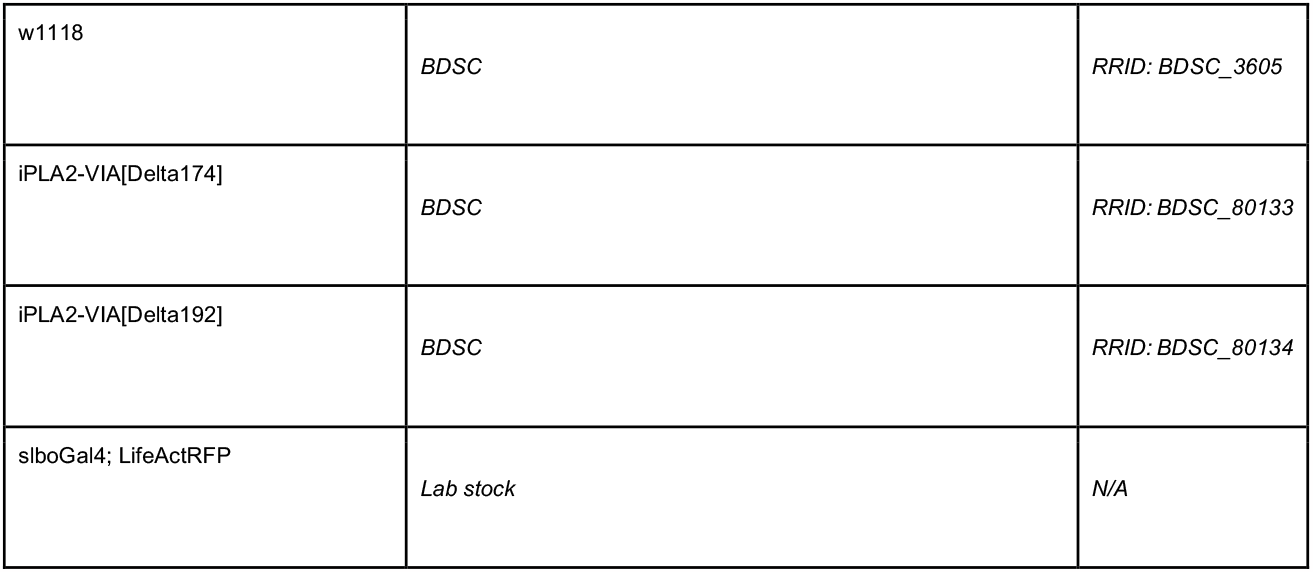
*Drosophila melanogaster* Stocks.

For Gal4 lines with Gal80^TS^, crosses were kept at 18 °C incubated at 29 °C for 3 days prior to dissection. For FlpOUT experiments to generate mosaic clones, flies were heat shocked 1 hour at 37 °C, restored to 25 °C for four hours, heat shocked again for 1 hour, and fattened for 3 days at 29 °C. Crosses with c306Gal4 were fattened for both 1 day and 3 days 29 °C prior to dissection for stage 10 as specified in Table 2. 2-5 day old fly progeny were supplemented with yeast for each experiment, and control and RNAi or overexpression flies in each experiment were matched in age. Table 2 reports genotypes and conditions for each analysis.

**Table 2.**
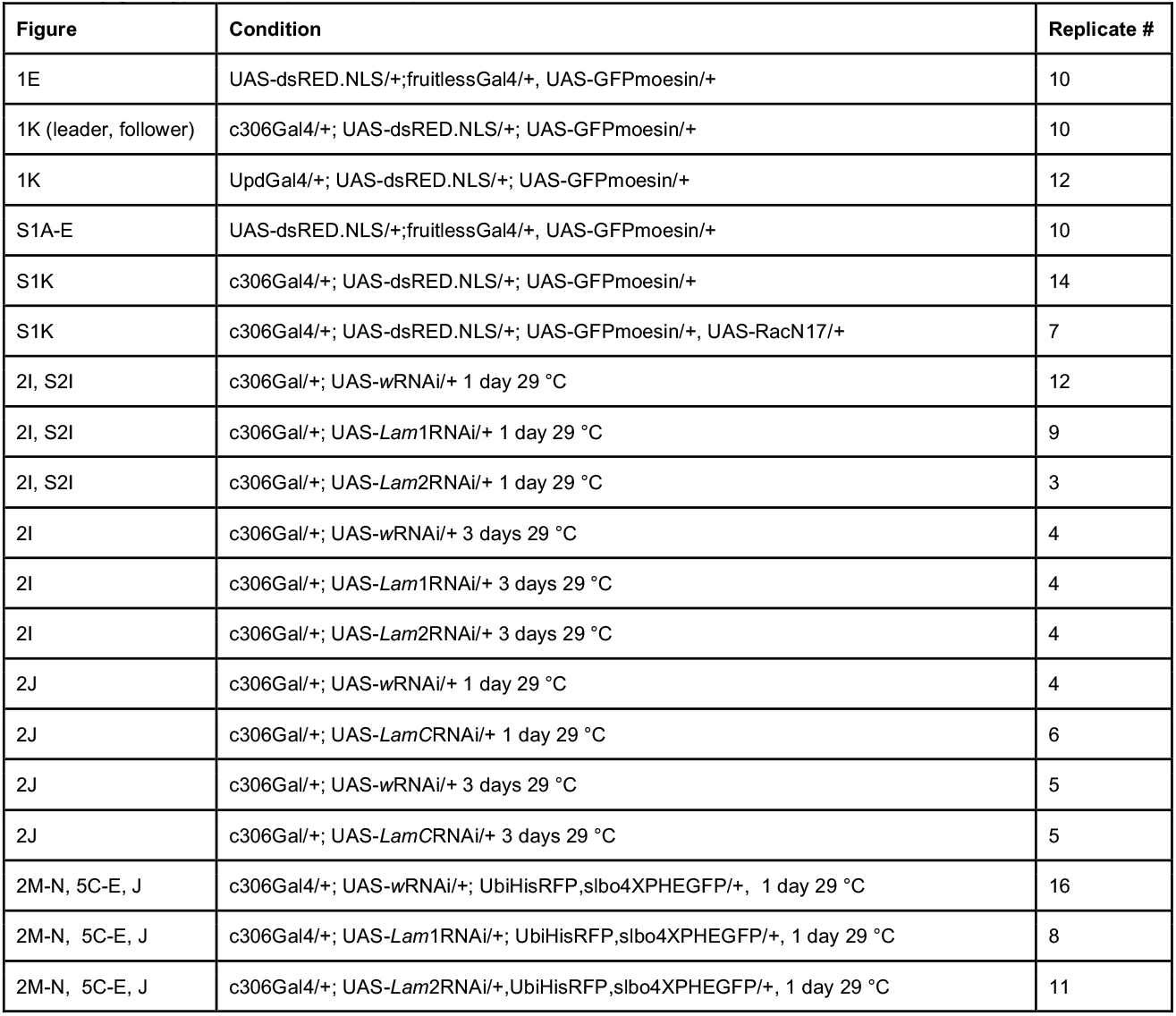
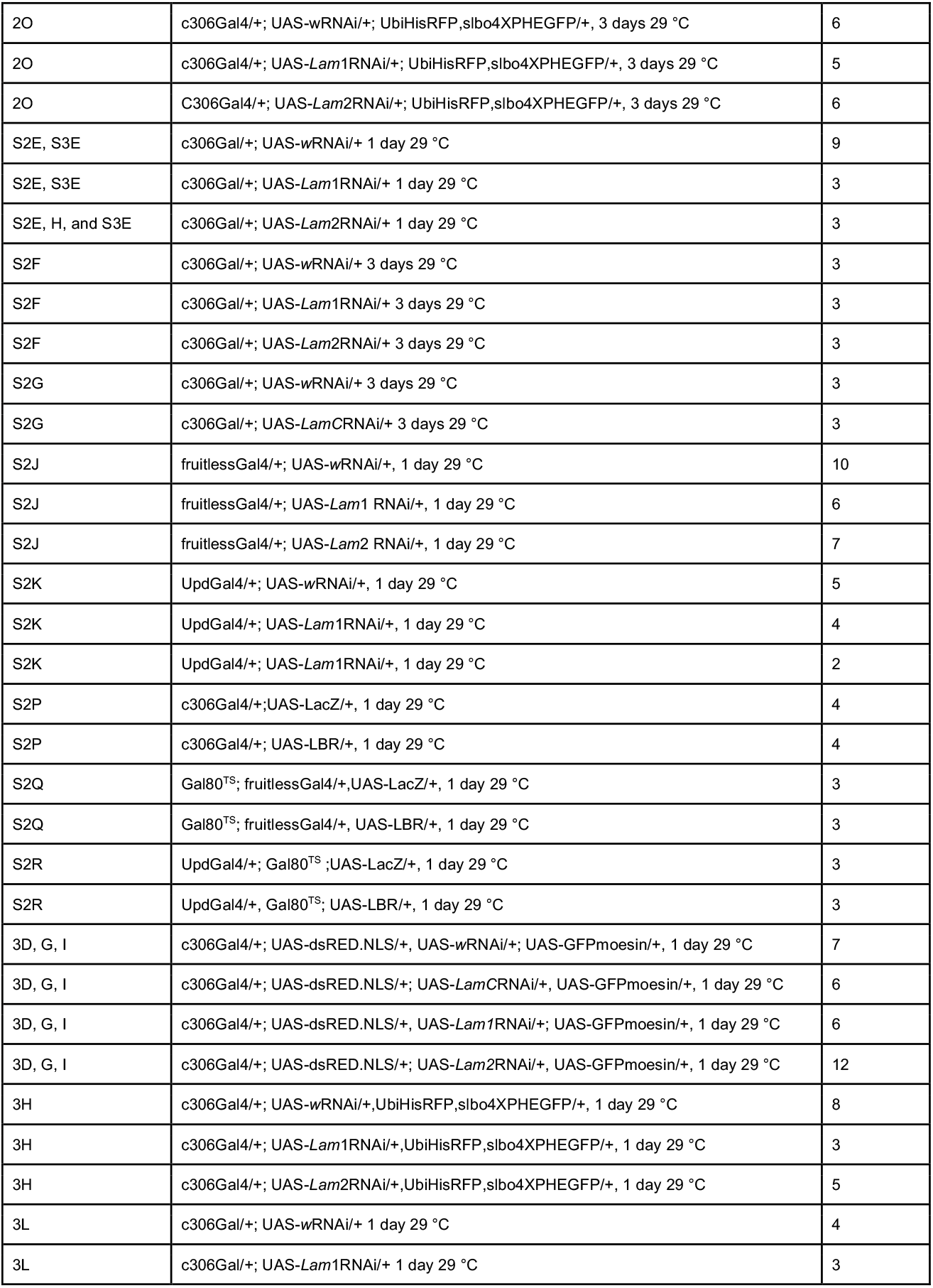
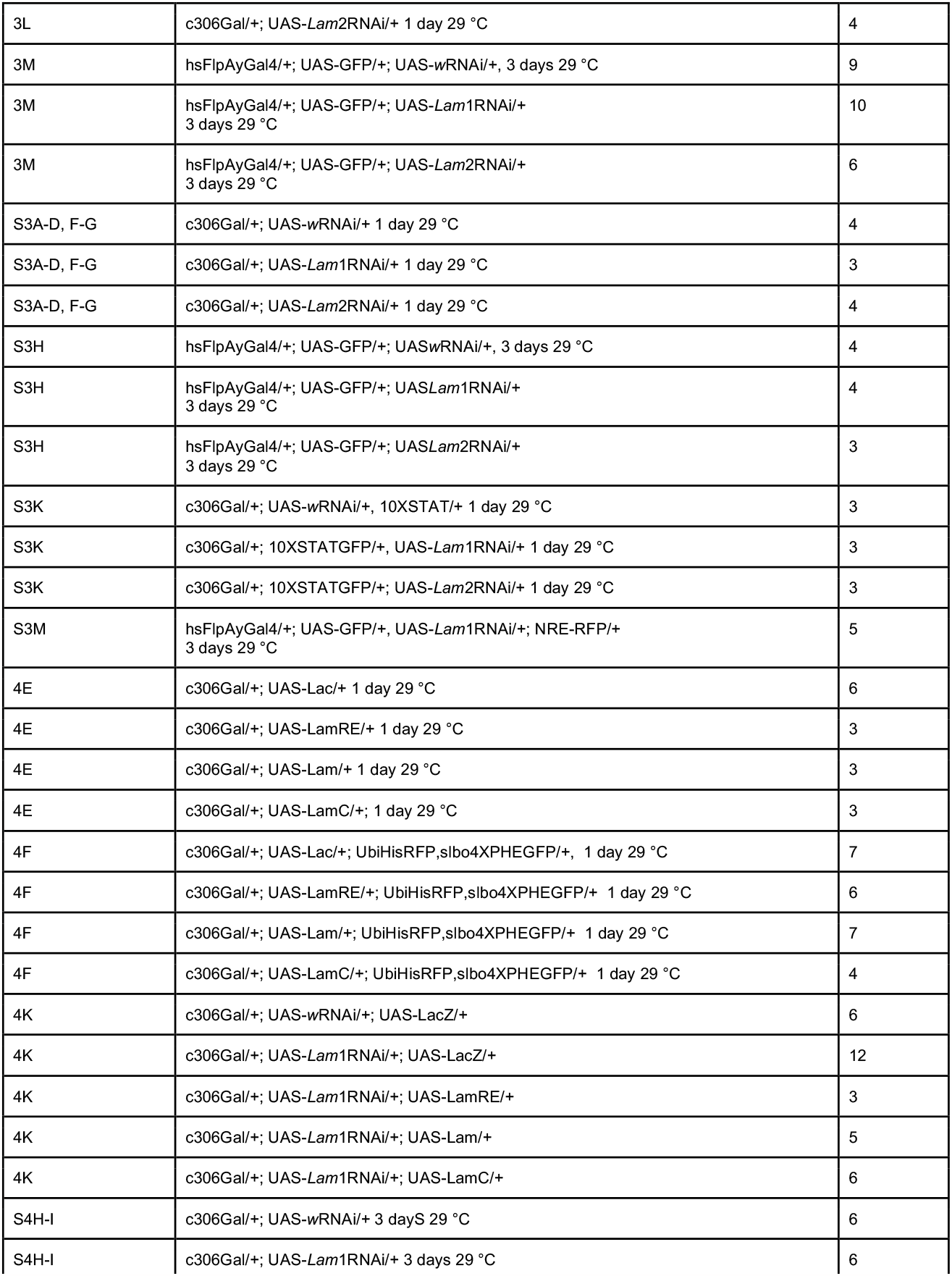
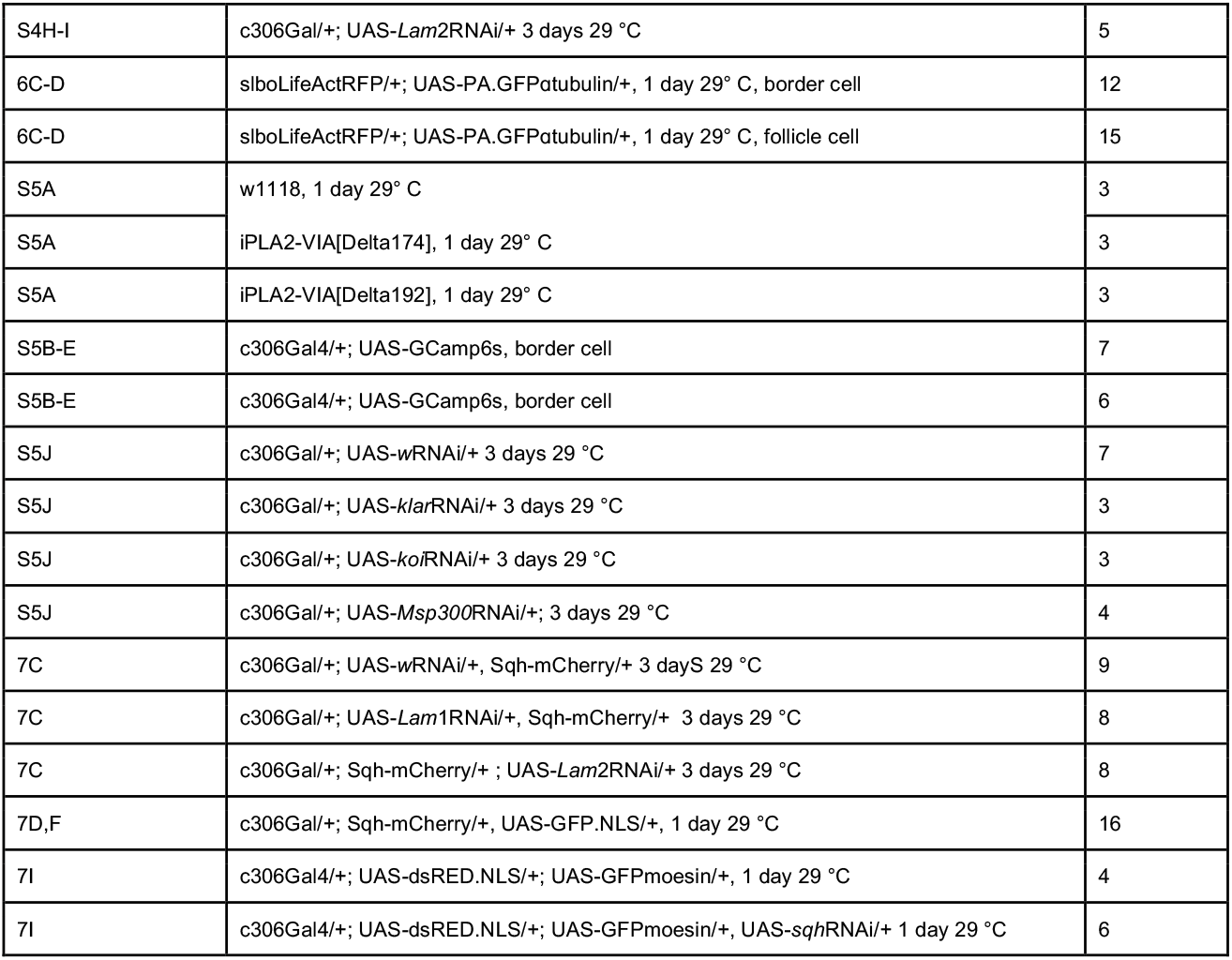
Fly genotypes and experimental replicates.

**Table 3.**
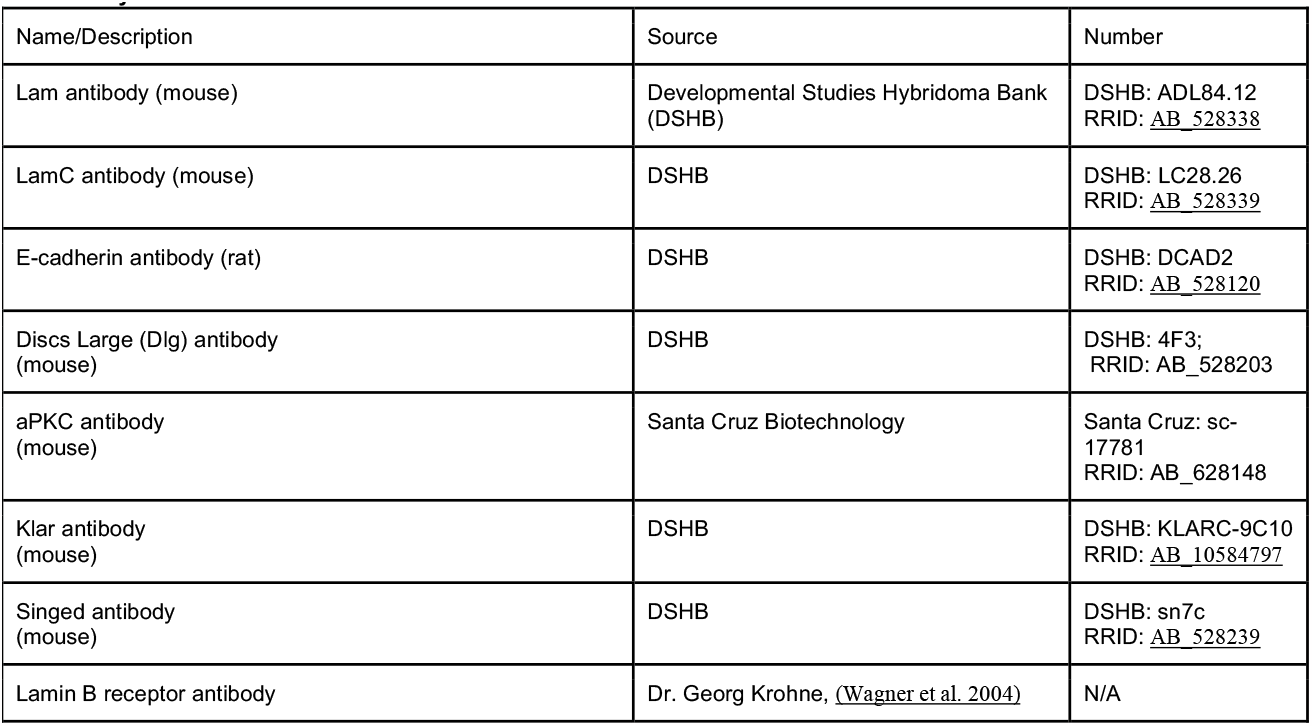
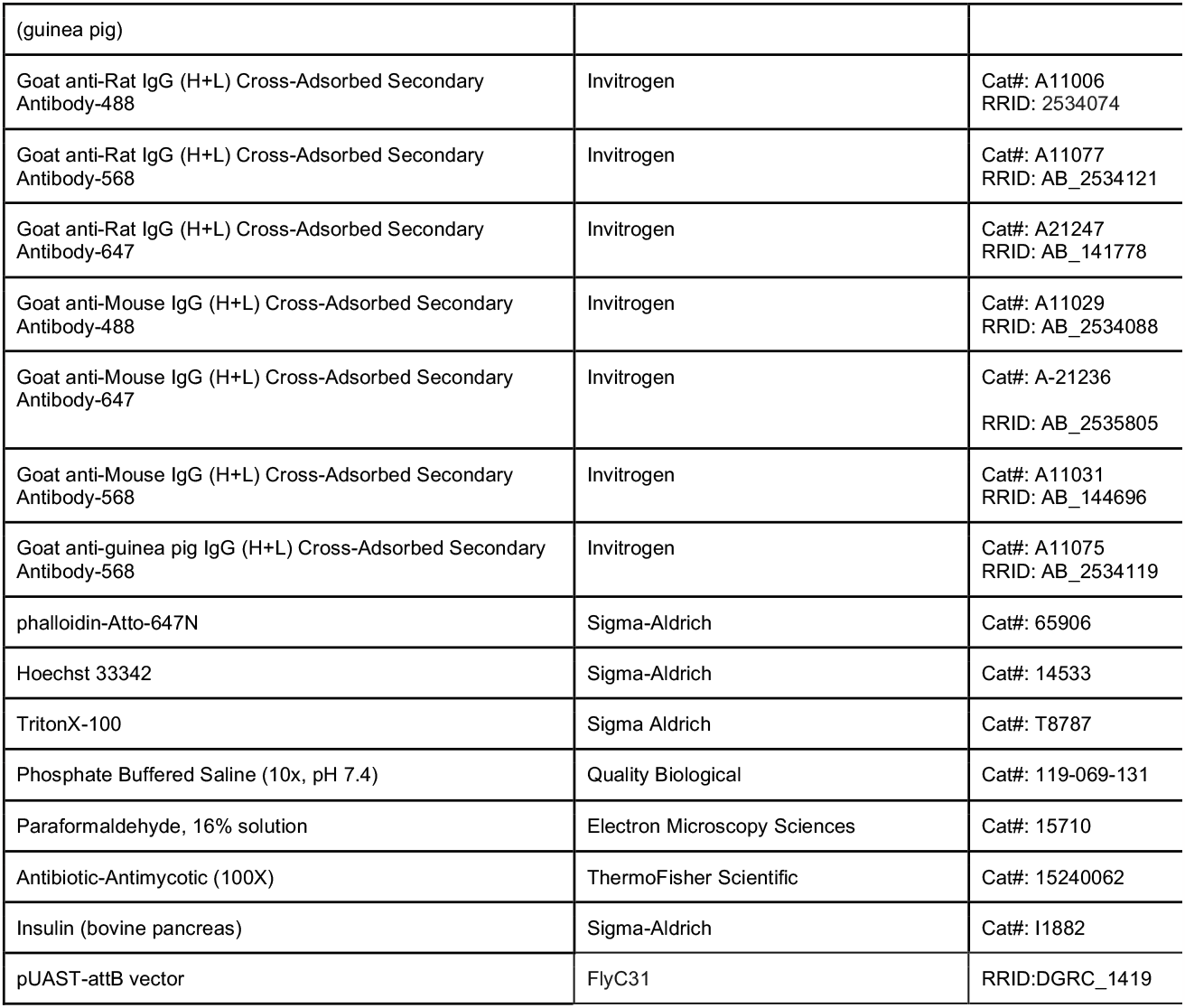
Key Antibodies and Chemicals.

### LamRE construct

The LamRE construct was generated to re-encode the DNA of the regions targeted by Lam1 and Lam2 RNAi lines in the coding sequence while keeping the same codon sequence. The re-encoded Lam coding sequence was ordered as a gene block(IDT, see coding sequence below). Then, it was introduced into a pUAST-attB vector(RRID:DGRC_1419) with in-fusion cloning (takara bio, #638947). The UAS-LamRE construct was sent to Bestgene to generate a transgene on chromosome 3 (stock RRID: BDSC_8622 (y^1^ w^67c23^; P{CaryP}attP2)) using phiCI integrase-mediated site-specific transgenesis.

### LamRESequence

ATGTCGAGCAAATCCCGACGTGCTGGCACCGCCACGCCGCAGCCCGGCAACACCTCCACCCCCCGGCCGCCATCGGCGGGTCCGCAGCCGCCGCCCCCCAGCACCCATAGCCAAACCGCTAGCTCCCCGCTGTCCCCGACGCGCCATAGCCGTGTCGCTGAAAAAGTCGAATTGCAAAATTTGAATGACCGTTTGGCTACGTATATCGATCGCGTCCGTAATTTGGAAACCGAAAATAGCCGTCTGACGATTGAAGTCCAAACGACGCGCGATACCGTGACCCGTGAAACGACGAATATTAAAAATATTTTTGAAGCTGAATTGTTGGAAACCCGTCGCTTGCTGGACGATACCGCCCGCGACCGTGCCCGCGCTGAAATTGACATTAAACGCCTGTGGGAAGAAAATGAAGAACTGAAAAATAAATTGGATAAAAAAACGAAAGAATGCACCACTGCTGAGGGCAATGTCCGCATGTACGAGTCGCGCGCCAACGAGCTGAACAACAAATACAACCAGGCCAACGCCGATCGGAAGAAGCTTAACGAAGACCTGAATGAGGCGCTAAAGGAGCTGGAGAGACTGCGTAAGCAGTTCGAGGAAACGCGGAAGAACCTGGAACAGGAGACACTGTCGCGCGTTGACCTGGAGAACACCATTCAGAGTCTGCGCGAGGAGCTCTCGTTCAAGGATCAGATCCATTCGCAGGAGATCAATGAGTCGCGCCGCATCAAACAGACAGAGTATAGCGAGATCGACGGTCGCCTCAGCTCCGAGTACGATGCCAAGTTGAAGCAGTCGCTGCAGGAGCTGCGCGCCCAGTACGAGGAGCAGATGCAGATTAATCGCGATGAAATCCAGTCCCTCTACGAGGACAAGATCCAACGACTGCAAGAGGCCGCCGCACGCACATCCAATTCCACGCACAAGTCCATCGAGGAGCTGCGCTCCACTCGTGTGCGTATCGATGCGCTCAACGCCAATATCAACGAACTGGAGCAAGCCAATGCCGACCTCAATGCGCGGATCCGTGATCTGGAGCGCCAGCTGGACAACGATCGCGAACGCCACGGTCAAGAGATAGACCTTCTCGAGAAGGAGCTCATTCGGCTGCGCGAAGAGATGACGCAACAGCTCAAGGAGTACCAGGACCTTATGGACATCAAGGTCTCCCTGGATTTGGAAATCGCCGCATACGACAAGCTGCTGGTGGGCGAGGAGGCTCGTTTGAACATCACCCCAGCCACCAACACGGCCACAGTGCAGTCCTTTAGCCAGTCGCTGCGCAACTCCACGCGAGCCACGCCATCGCGTCGCACTCCCTCTGCTGCCGTGAAGCGCAAACGCGCCGTGGTCGACGAGTCGGAGGATCACAGCGTCGCCGATTACTATGTGTCCGCCAGTGCCAAGGGCAACGTGGAGATCAAGGAGATCGATCCCGAGGGCAAGTTCGTAAGGCTGTTCAACAAGGGCAGCGAGGAGGTGGCCATCGGTGGCTGGCAGCTGCAACGCCTGATTAATGAAAAGGGCCCCAGCACGACCTATAAATTTCACCGCAGCGTCCGCATTGAACCCAACGGAGTCATTACGGTGTGGAGCGCCGATACGAAAGCTAGCCATGAACCCCCCAGCTCCCTGGTCATGAAAAGCCAAAAATGGGTGAGCGCTGATAATACCCGCACCATCCTGTTGAATAGCGAAGGAGAAGCTGTCGCTAACTTGGACCGTATTAAACGTATCGTCAGCCAGCATACCAGCAGCAGCCGCTTGAGCCGCCGCCGTTCCGTCACGGCTGTCGATGGAAACGAACAACTGTATCATCAACAAGGAGACCCCCAACAAAGCAATGAAAAATGTGCTATCATGTAG

### Immunostaining and Fixed Imaging

Ovaries were dissected in Schneider’s media (Thermo Fisher Scientific) supplemented with 20% heat-inactivated Fetal Bovine Serum (Sigma-Aldrich) and 1x antimycotic/antibiotic (VWR) and adjusted to a pH of 6.85-6.95. Ovaries were then fixed in 4% paraformaldehyde in 1X Phosphate-buffered saline (PBS) for 15 minutes. Ovarioles were washed 3x 10 minutes with 1X PBS +0.4% triton (PBST) and incubated in primary antibodies overnight at 4 °C. Antibodies used include: Lam (1:10, Developmental Studies Hybridoma Bank(DSHB), #ADL84.12), E-cadherin (1:15 DSHB #DCAD2), LamC (1:20, DSHB #LC28.26), singed (1:20, DSHB, #sn 7c), Klar (1:10 DSHB, #KLARC-9C10), Lamin B receptor (1:200 gift from Dr. Georg Krohne), aPKC (1:500, Santa Cruz Technologies), Dlg (1:20, DSHB, #4F3). The following day, ovaries were washed with 1X PBST followed by 3×10 minutes washes in 1X PBST. Then, ovaries were incubated with secondary antibodies (Goat anti-Rat IgG (H+L) Cross-Adsorbed Secondary Antibody, Alexa Fluor 647, 488, or 568, Invitrogen, Goat anti-Mouse IgG (H+L) Cross-Adsorbed Secondary Antibody, Alexa Fluor 488 and 568, Invitrogen), phalloidin-Atto-647N (1:500, Sigma-Aldrich), and Hoechst (1:500, Invitrogen) for 1 hour at room temperature. Ovaries were washed with 1 X PBST followed by 3×10 minutes washes in 1X PBST. After the last wash, ovaries are suspended in vectashield (H-1000, Vector Laboratories) and incubated overnight at 4 °C before mounting onto slides. Details for reagents and sources are listed in Table 3.

Slides were imaged on a Zeiss LSM780 confocal microscope with a 40× 1.1 N.A. 0.62mm long working distance water objective.

### Live Imaging

Ovarioles were dissected from ovaries in live imaging media (Schneider’s media (Thermo Fisher Scientific) supplemented with 20% heat-inactivated Fetal Bovine Serum (Sigma-Aldrich) and 1x antimycotic/antibiotic (VWR) with a pH of 6.85-6.95) and resuspended in live imaging media with 0.4 mg/mL bovine insulin (I1882, Sigma-Aldrich) and 1% agarose and immediately mounted for imaging. Stage 9 egg chambers were imaged on an inverted Zeiss LSM800 confocal microscope fitted with a Plan-Apochromat 40X, 1.2 NA multi-immersion objective with the collar set for water immersion at 3, 2 minutes or 20 seconds time intervals as specified in figures and movies.

For imaging with dextrans to mark extracellular spaces, fluorescent dextrans were added to the media prior to mounting (100 ***μ***g/ml; Dextran Alexa 647 10,000 MW; Thermo Fisher Scientific; D22914).

### Image analysis

#### Diameter measurements

For each movie, the diameters of clusters, cells, nuclei, and junctures in the central path were measured prior to delamination for the first timeframe (Fig. 1E). The outline of each cell, nucleus, and the entire cluster were manually traced from fluorescent markers in ImageJ (GFP-moesin for border cells, and dsRED.NLS for nuclei and dextran-Alexa647 junctures).

### Protrusion shape analysis

The time of delamination (Fig. 1F, S1A-B) was scored as the full detachment of all border cells from the anterior of the egg chamber and set as time=0 for the measurements and all time points were set relative to delamination=0 minutes. UAS-GFP-moesin actin binding domain was expressed in border cells by fruitlessGal4 or slbo4XPHEGFP was used to mark the border cells boundaries. To measure protrusion length, a line that was parallel to the long axis of the egg chamber was drawn from the polar cell boundary to the protrusion tip. To measure protrusion width, lines were manually drawn across the middle of the protrusion neck to the long axis of the egg chamber (for Fig. S1A Fig. 5C-E) or at the tip or base of the protrusion (Fig. S1C-E, Fig. 5F-J) and measured in Image J for each time point.

### Nuclear movement and shape analysis

To measure nuclear position in the protruding cells, a line was manually drawn from the back of the nucleus to the nearest polar cell membrane marked with slbo4XPHE-GFP for each time point. The change in nuclear position was measured as the change in length of the line between two time points and divided by the time interval. Only the largest protrusion was scored for each timepoint if there were multiple protrusions, and analysis was only performed in protruding cells. To set a threshold for forward, neutral or backward nuclear movement, the average and standard deviation were calculated for the data set, and forward movement was defined as >1 standard deviation (SD) greater than the mean, neutral was defined as within 1 SD from the mean, and backward was defined as more than 1 SD lower than the mean.

To measure the lead cell’s nuclear aspect ratio relative to the migration axis lines (Fig S1A-B, 3H), lines were manually drawn across the center of the nucleus marked with dsRED.NLS parallel and perpendicular to the long axis of the egg chamber. The aspect ratio was calculated by dividing the parallel nuclear diameter by the perpendicular nuclear diameter.

To measure the nuclear aspect ratio and circularity (regardless of egg chamber orientation) from time-lapse series for comparison of leader, follower, and polar cell nuclei, the central z-slice of the channel with UAS.dsRED.NLS in each nucleus was cropped from each time point to prevent overlap of nuclei during semiautomated analysis. Nuclei were segmented into binary images using a custom Matlab code (available upon request), and shape parameters were measured over time using the analyze particle tools in ImageJ. We manually verified if the segmentation worked properly and whether the nucleus was in view of not. If the automated analysis did not properly segment the nucleus in a time point, a manual trace was performed.

Samples were blinded throughout the analysis. All timeframes were included unless the nucleus went out of focus during the imaging session. The average nuclear aspect ratio was automatically generated in ImageJ measurements. The change in nuclear aspect ratio was calculated by taking the absolute value between two aspect ratios of adjacent time points.

For fixed imaging analysis of nuclear shape, the Hoecsht signal in fixed border cells proved difficult to segment automatically. Therefore, each nucleus in a border cell cluster was traced manually in Image J by using the Hoechst signal to mark the nucleus in stage 9 egg chambers. All border cells’ nuclei were included unless Hoechst staining did not allow distinguishing of a nucleus from background or if a nucleus was out of focus. ImageJ measurements were used to calculate area and aspect ratio of traces of nuclei in fixed images.

### Myosin II flash analysis

Each movie was aligned so that the X axis was the long axis of the egg chamber, and the anterior was on the left. The egg chamber was aligned over timepoints to correct for drift using the StackReg plugin in ImageJ. Myosin II flashes were manually scored for each time point relative to their position of the nucleus aligned to the migration axis (front: further from than nucleus, side: parallel in X axis to nucleus, back: closer to anterior than nucleus) It was also noted if there was no flash near the nucleus at the timepoint. Nuclear movement defined as X axis displacement was measured by tracking the nuclear position over time using manual tracking. Nuclear shape measurements with the GFP.NLS was measured using the same semi-automatic analysis method described for dsRED.NLS above. Then, the average movement and shape change were binned based on the presence and position of flash for each movie.

### Border cell size

To measure border cell size, cytoplasmic GFP in FlpOut clones expressing wRNAi or LamRNAi were automatically segmented using a Matlab code and analyzed for area in ImageJ using the analyze particles. Only clones that were distinguishable from other clones in a cluster were used for segmentation.

### Border cell migration indexes

To measure border cell migration defects, all stage 10 egg chambers were identified by onset of centripetal cell migration, and the border cell cluster was detected with E-cadherin or phalloidin staining. The position of the border cell cluster was manually scored based on positions (0%: still attached, <50% migrated relative to distance between anterior and oocyte boundary, >50% migrated relative to anterior:oocyte, and 100%: at oocyte boundary). Then, the position of the cluster was recorded for each stage egg chamber. RNAi and overexpression experiments were performed at least 3 times independently with 3 independent genetic crosses, and with the control line (UAS-whiteRNAi or UAS-LacZ) crossed in parallel.

### Migration speed and Directionality Index

To measure the average speed of each border cell cluster, the z-slices from time-lapse series were max-projected. Then, the egg chamber was aligned over timepoints to correct for drift using the StackReg plugin in ImageJ. The manual tracking plugin in ImageJ was used to track border cell cluster position over time.

The directionality index was measured as previously reported (Cai et al., 2014). The net distance traveled over the time lapse movie was divided by the sum of the distances moved in each time point.

### Lamin staining and border cell number

In stage 10 egg chambers, the number of border cells was manually counted using Hoechst and border cell markers (E-cadherin or Phalloidin). These cell number counts were limited to 30 micron z-sections, which usually covered the entire cluster. Lam or LamC depletion was categorized as “none” (complete rim around the Hoechst signal), “partial” (incomplete rim around Hoechst), or “full” (no detectable lamin at the nuclear rim). The percent of border cells with full, partial, or no knockdown were divided by the total number of border cells in the cluster. The averages from each LamRNAi line are shown in Fig. S2E-G. Then, the averages were calculated for completed migration or incomplete migration in the Lam2RNAi (line #45635) (Fig. S2H)

### STAT reporter quantification

To measure STAT activity in Stage 8 egg chambers, we crossed c306Gal4/Y; 10XSTATGFP (STAT92e)/CyO to UAS-LamRNAi or UAS-wRNAi lines. We performed immunostaining and imaged the anterior of stage 8 egg chambers. All egg chambers were imaged using the same conditions. For each egg chamber, we drew a small region of interest in each of the 2 cells adjacent to the polar cells in ImageJ and acquired the average fluorescent intensity and then subtracted a box in the nurse cells as background. Then, averages were normalized to the average of the UAS-wRNAi control to show the relative change in the fluorescence of the STAT reporter from the control.

### Notch Responsive element quantification

To measure Notch activity, we generated a fly co-expressing UAS-LamRNAi (line #107419) or UAS-wRNAi with a Notch Responsive Element-Red Reporter (NRE-RR) (Zacharioudaki and Bray, 2014), and crossed these flies to a hsFlpAyGal4;UAS-GFP line. We heat shocked progeny as described above to generate clones. Only clusters with GFP+ clones were used for analysis. For measurements of NRE-RR fluorescent intensity, a box overlaying the nucleus was drawn in the center of each border cell and subtracted the average fluorescent intensity of a box drawn in the nurse cells (background). For each cluster, the average fluorescence intensity of the GFP+ clones was divided by the average fluorescent intensity of the GFP-clones in the same cluster.

### E-cadherin and F-actin quantification

For the total mean levels, each border cell cluster still juxtaposed to the anterior was traced and the mean fluorescence intensity was measured for F-actin and E-cadherin Circularity of the cluster was measured from these same traces of the cluster.

For the peripheral E-cadherin analysis, FlpOUT clones with WT and LamRNAi clones were analyzed. The mean of 3 pixel thick lined traced around the periphery of GFP+ clone (UAS-LamRNAi) was divided mean periphery of a WT clone in the same cluster.

For the distribution of F-actin, a 10 pixel thick line scan was drawn across the cluster from front to back and the levels were measured. Lines were normalized and binned to a position where the top 10% of the line was deemed the front, the 11-89% was deemed the middle, and the last 10% was deemed the back for each trace.

### GCaMP Imaging and Quantification

GCaMP signal was monitored every 5 seconds over a total of 5 minutes as described previously to monitor CaMP flashes(Sahu et al., 2017). A region of interest was drawn in one z-plane at the front of the border cell cluster and the average fluorescence intensity (F.I.) measured over time. As a positive control, a region of interest was drawn in a follicle cell displaying GCaMP flashes. The average fluorescence intensity and the maximum fold change (F.I._time1-F.I_.time2)/F.L._time1 was measured from those values.

### Photo-activation experiments

For photoactivation of GFP-αtubulin, a region of interest (ROI) was drawn in the protrusion of a leading cell or on one side of the nucleus in a posterior follicle cell. Photo-activation was performed on Zeiss 800 with 405 laser at 20% power for 3 iterations every 2 seconds. Photoactivation started after 6 seconds and all movies were captured over 120 seconds total. Then, a back and front ROI were drawn in image day and the mean was measured for each time point. Data were normalized to the front:back ratio of the first time point for Figure 6C. To measure the time constant, we fit each curve of the mean intensity over time to a Boltzmann sigmoidal function as done previously (Petrie et al., 2014) using PRISM. The slope of the curve is the time constant and inversely proportional to diffusion. All slopes were included in the figure despite the R_2_ value, but removing data that did not fit the curve R^2^< 0.8 did not change results.

### Statistical analysis and data presentation

Data points, middle bars and error bars are specified in the legends but generally refer to the mean +/-SEM with each data point plotted. Experimental replicates are reported in Table 2 and number of biological replicates (# of egg chambers) are specified in figures and legends. Inclusion criteria: all undamaged samples and in focus images were included for analysis. Exclusion criteria: damaged samples and out of focus timepoints and images. Nuclear shape analyses from live imaging were blinded. All other measurements were not blinded but performed in a systematic manner to minimize bias. Power calculations were not generally performed, but we aimed to perform all experiments in triplicate to have sufficient power for statistical testing. Genetic crosses were performed at least in duplicate. Flies that were established genetic crosses and flies used for experiments were selected randomly from vials of the appropriate age and genotype, but there were no procedures that required randomization of subjects. Data were analyzed for normality using a Shapiro-Wilk normality test in Prism. A t-test or a one way ANOVA (parametric test) was performed if data passed the normality test. If data did not pass the normality test, a nonparametric test was performed in prism. Statistical tests used and sample size are reported in figure legends. Fluorescence images were acquired and processed exactly the same between control and experimental conditions. Figure legends include additional information on data representation for each specific experiment.

### Online supplementary material

Fig. S1 shows nuclear shape dynamics relative to protrusion and cluster dynamics in border cells. Fig. S2 shows LamC and Lam are depleted with RNAi lines, Lam is required in outer border cells, and LBR overexpression is required for nuclear shape and for migration in outer border cells. Fig. S3. shows border cell organization and gene expression in Lam-depleted clusters versus controls. Fig. S4 shows border cell polarity in control and Lam-depleted clusters. Fig. S5. shows iPLA2 mutants and clusters depleted of LINC complex components complete migration and shows examples of Myosin II and calcium dynamics in border cell clusters. Video 1. shows border cell delamination. Video 2. shows leading nuclei deform rapidly. Video 3. shows leader cells undergo the most deformation while polar cells do not have major shape changes. Video 4. shows control clusters delaminate and form a main protrusion while Lam-depleted clusters fail to invade and stabilize a protrusion. Video 5. shows protrusion expansion in controls versus protrusion narrowing in Lam-depleted clusters relative to nuclear dynamics. Video 6. Shows photoactivation of PA-GFP-tubulin in a leading border cell and follicle cell. Video 7. shows GCaMP in border cells and follicle cells. Video 8 shows Myosin II dynamics in control and Lam-depleted clusters. Video 9. shows myosin II dynamics correlate with nuclear dynamics. Video 10 shows Myosin II depleted results in failed delamination, long lived protrusions, and lack of nuclear movement. Table 1. lists *Drosophila melanogaster* stocks. Table 2. reports genotypes, experimental conditions, and replicates for figures. Table 3 reports key antibodies and chemical reagents.

## Supporting information

Video 1. Border cell cluster undergoing delamination.

Video 8. Myosin II dynamics in control and Lam-depleted clusters

Video 9. Myosin II flashes correspond with nuclear movement.

Video 2. The nucleus in the leading cell undergoes rapid changes in shape.

Video 10. Myosin II depletion results in long-lived protrusions and little nuclear movement.

Video 7. Calcium dynamics in border cells.

Video 6. Photoactivation of GFP-alpha tubulin

Video 5. Nuclear stretching and movement corresponds with protrusion widening while Lam-depleted cells do not maintain protrusions.

Video 4. Control border cells maintain a leading protrusion as they delaminate and migrate while Lam-depleted clusters form ectopic protrusions and ha

Video 3. Border cell nuclei undergo dynamic shape changes while polar cells maintain nuclear shape.

## End Matter

### Data Availability

Data are shown in the figures and supplemental files, and additional data files will be posted on a public repository (Figshare) or made available upon request.

## Acknowledgments

We would like to thank the Bloomington Drosophila Stock Center, the Vienna Drosophila Resource Center (VDRC) for the Drosophila stocks. We would like to thank Georg Krohne for the Lamin B receptor antibody, Lori Wallrath for the UAS-LamC fly stock, and Jörg Großhans for the UAS-Lam fly stock. We would like to thank Carinna Tran, Emory Campbell, and Dr. Jim Mondo for their technical assistance, Dr. Joseph Campanale and Dr. Maddalena Nano for their careful reading of the manuscript and all of the members of the Montell lab for their feedback on this work.

## Author Contributions

L.P. and D.J.M. designed research, L.P. performed research, L.P. analyzed data; and L.P and D.J.M wrote the paper.

---

**Figure S1.**
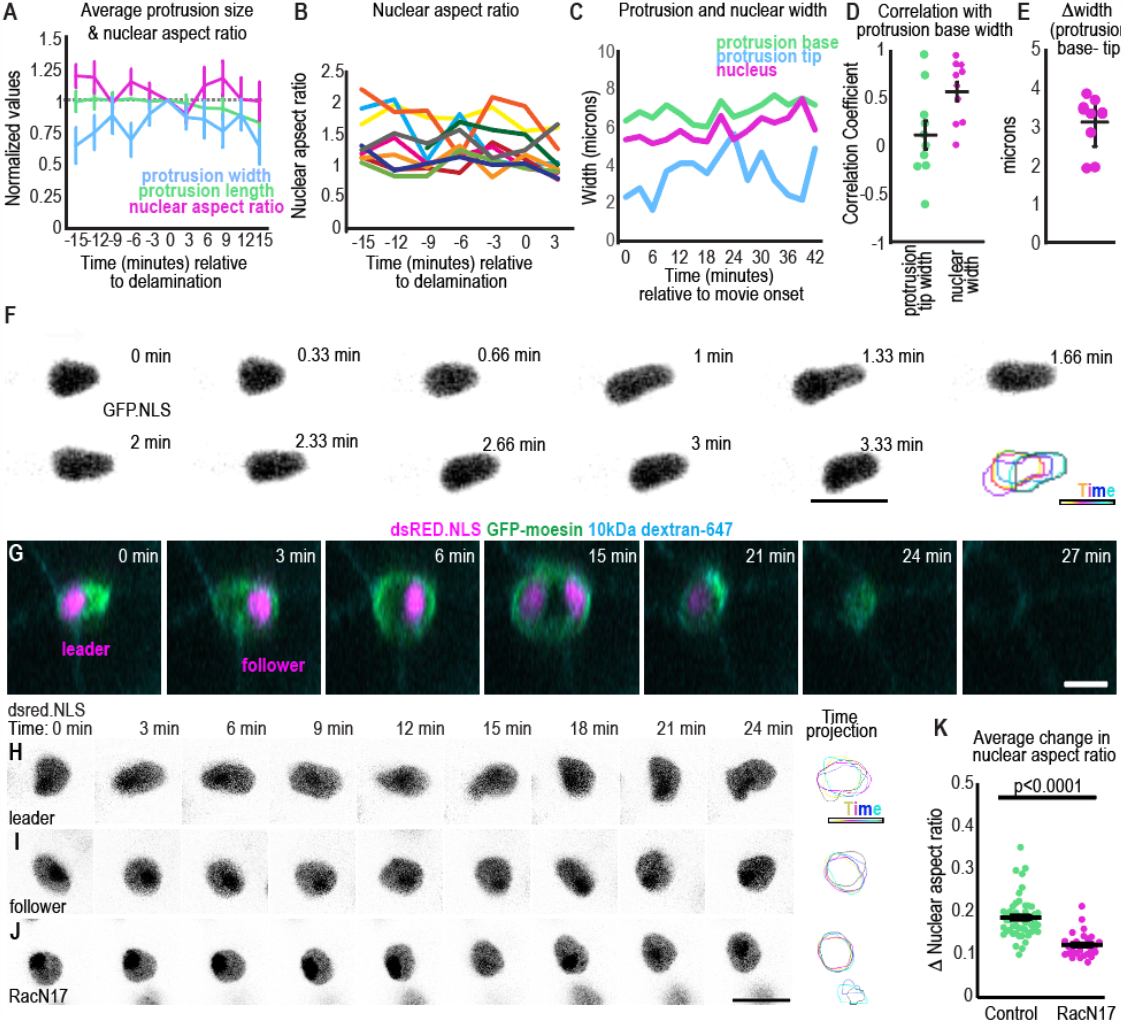
Changes in nuclear shape occur rapidly in migrating border cell clusters. A. Plots of the average +/- SEM of each indicated parameter relative to the time of delamination. N=10 movies. B. Nuclear aspect ratio of each leading cell’s nucleus aligned to the protrusion axis. C. Representative plot showing example where nuclear width and protrusion width simultaneously oscillate while the protrusion tip width changes variably. D. Average correlation between nuclear width and protrusion base width or between protrusion tip and base width. Unpaired t-test p=0.02. E. Average difference between the protrusion base and protrusion tip. F. Images from a timelapse series with the leading cell nucleus marked with GFP.NLS. Images were acquired every 20 seconds. Bottom right: projection of timepoints.G. ZY view of border cell cluster leading cell and following cell over time to show the expansion and shrinking of the juncture as the cluster moves through it. H-J. Example images of nuclei from time-lapse series of control leader, follower or RacN17 dominant negative expressing border cells. Right: projection of timepoints. K. Plot of the average change in nuclear aspect ratio, bars: mean +/- SEM dots: value for one nucleus. A Mann-Whitney test was used to test for statistical significance. Scale bars: 10 ***μ***m. Genotypes and experimental replicates reported in Table 2.

**Figure S2.**
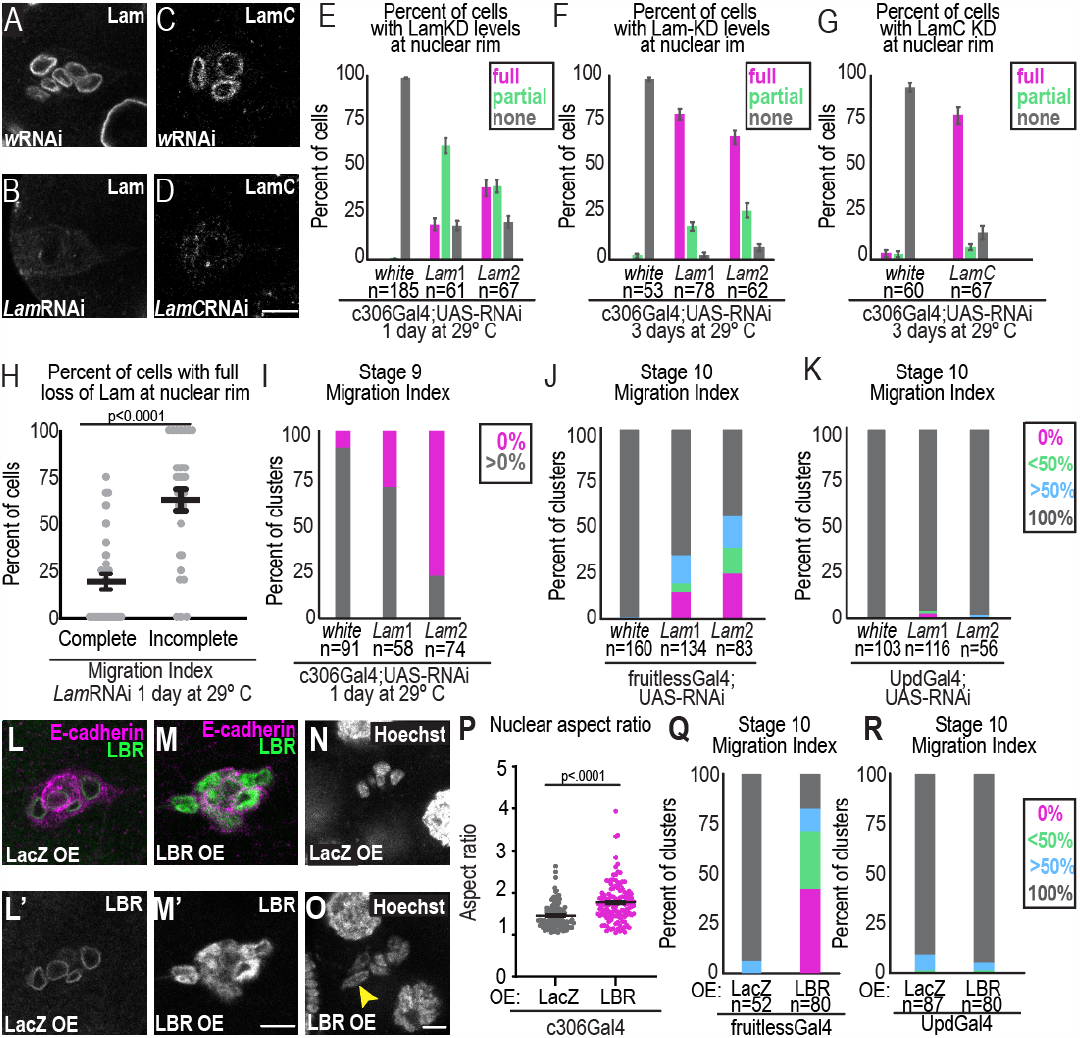
Stronger depletion of lamins correlates with higher rates of migration defects. A-D. Images of Lam (A-B) and LamC (C-D) for the indicated conditions. E-G. Plots showing the mean +/- SEM percentage of border cells with the indicated depletion (full, partial or none) of Lam or LamC at the nuclear periphery for the indicated conditions. E. A Kruskal Wallis Test was performed to compare means. mean with the full KD: wRNAi versus Lam1, p=0.006 and w versus Lam2 p<0.0001. F. Kruskal Wallis Test was performed to compare those with a full KD: w versus Lam1, p<0.0001 and w versus Lam2 p<0.0001. G. Kruskal Wallis Test was performed to compare the mean of cells with a full KD, wRNAi versus LamC p<0.0001. H. Plot showing the individual and average percent of cells with indicated knockdown in Lam-depleted clusters with complete or incomplete migration. A Kruskal-Wallis test was performed. Data shown is from line #Lam2RNAi after 1 day at 29 ºC (N=3 experimental replicates). I. Migration Index of stage 9 egg chambers for the indicated conditions to quantify the percent that have moved away from the anterior (grey). n=number of egg chambers, after 1 day at 29 ºC. A Fisher’s exact test with a Bonferroni correction to compare % with delamination yield p=0.0003 for wRNAi versus Lam1RNAi, and p<0.0002 for wRNAi versus Lam2RNAi. J-K. Migration Index for the indicated conditions, n= number of egg chambers. A Fisher’s exact test with a Bonferroni correction to compare % with complete migration (p values for J: wRNAi versus Lam1RNAi p<0.0002, w RNAi versus Lam2RNAi p<0.0002) (p values for K: wRNAi versus Lam1RNAi, p=0.4 wRNAi versus Lam2RNAi, p>0.9). KD condition: 1 day at 29 ºC. L-M Images of fixed border cells from stage 9 egg chambers stained for Lamin B receptor (LBR) with LacZ overexpressed (L) LBR overexpressed (M) with c306Gal4 for 1 day at 29 ºC. N-O. Images of Hoechst (DNA) staining of stage 9 egg chambers overexpressing LacZ(N) or LBR(O) kept at 29 ºC for 1 day. Scale bars: 10 ***μ***m. P. Plots measuring nuclear shape in nuclear aspect ratio from Hoechst channel of fixed border cell. Each dot represents an individual nucleus and middle and error bars represent the mean +/- SEM. A Mann-Whitney test was used for statistical testing. N=3 experimental replicates. Q-R. Plots of migration indexes of staged egg chambers in the noted conditions. n=number of egg chambers counted. A Fisher’s exact test was used to compare % with complete migration and yields p<0.0001 for Q. and p=0.5 for R. Genotypes and experimental replicates reported in Table 2.

**Figure S3.**
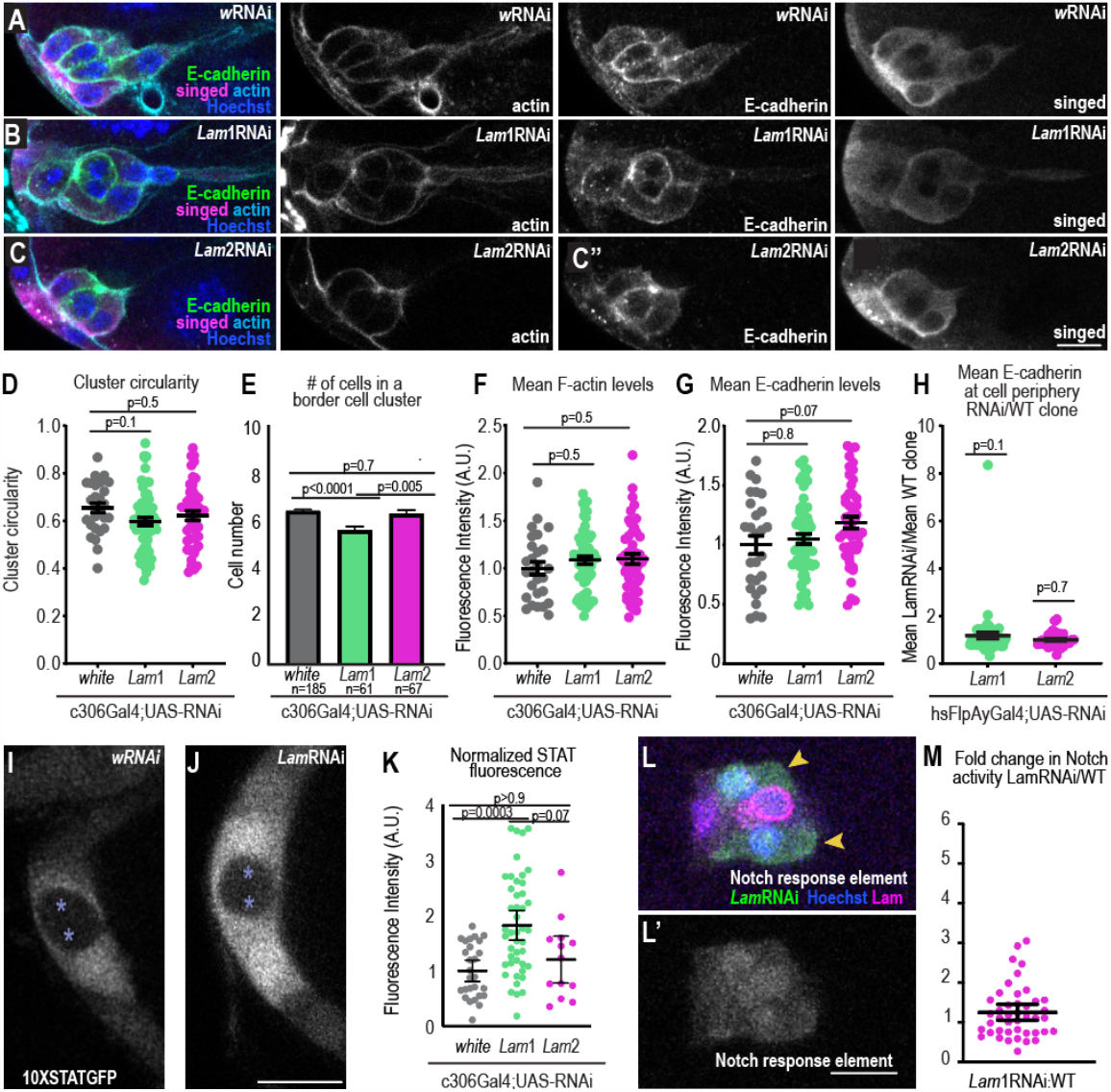
Lam-depleted border cells specify and express key border cell genes. A-C Representative images of stage 9 border cells stained with the indicated markers in control (A) and LamRNAi (B) and Lam2RNAi (C). Left: merged images. Right: grayscale images of F-actin, E-cadherin, and singed staining. D. Plot of individual (dots) and average +/- SEM(bars) cluster circularity, 1 day at 29 ºC. An ANOVA with Tukey post-hoc was performed. E Plot of average +/- SEM numbers of border cells in a cluster, 1 day at 29 ºC. A Krusal-Wallis test was performed. F-G Mean F-actin and E-cadherin levels for each clusters (dots) and the average +/- SEM (bars). KD condition: 1 day at 29 ºC (see also Fig. S5 for 3 day F-actin analysis). An ANOVA with Tukey post-hoc was performed for each plot. H. Clonal analysis the ratio of mean peripheral E-cadherin in a LamRNAi clone divided by the mean of a WT clone. KD condition: 3 days at 29 ºC. Middle bars show the mean +/- SEM. A Wilcoxon test was performed. I-J Confocal images of STAT activity reporter (10XSTATGFP) for indicated conditions. LamRNAi line shown: Lam1RNAi, KD condition: 1 day at 29 ºC. K. Plot showing individual (dots) and mean +/- SEM (bars) measures of STAT fluorescence normalized to the mean of the control. A Kruskal-Wallis test was used for statistical testing. L. A border cell cluster expressing hsFlpAyGal4 UAS-GFP,LamRNAi clones and wildtype clones with the indicated markers a merge of channels. L’ Greyscale image of notch responsive element-RFP. M. Plot showing the fluorescence intensity of the nuclear Notch responsive element relative to wildtype clones in the cluster, KD condition: 3 days at 29 ºC. Middle bars show the mean +/- SEM. A Wilcoxon test was used to test for statistical upregulation, p=0.06, ns. Scale bars: 10 ***μ***m. Genotypes and experimental replicates reported in Table 2.

**Figure S4.**
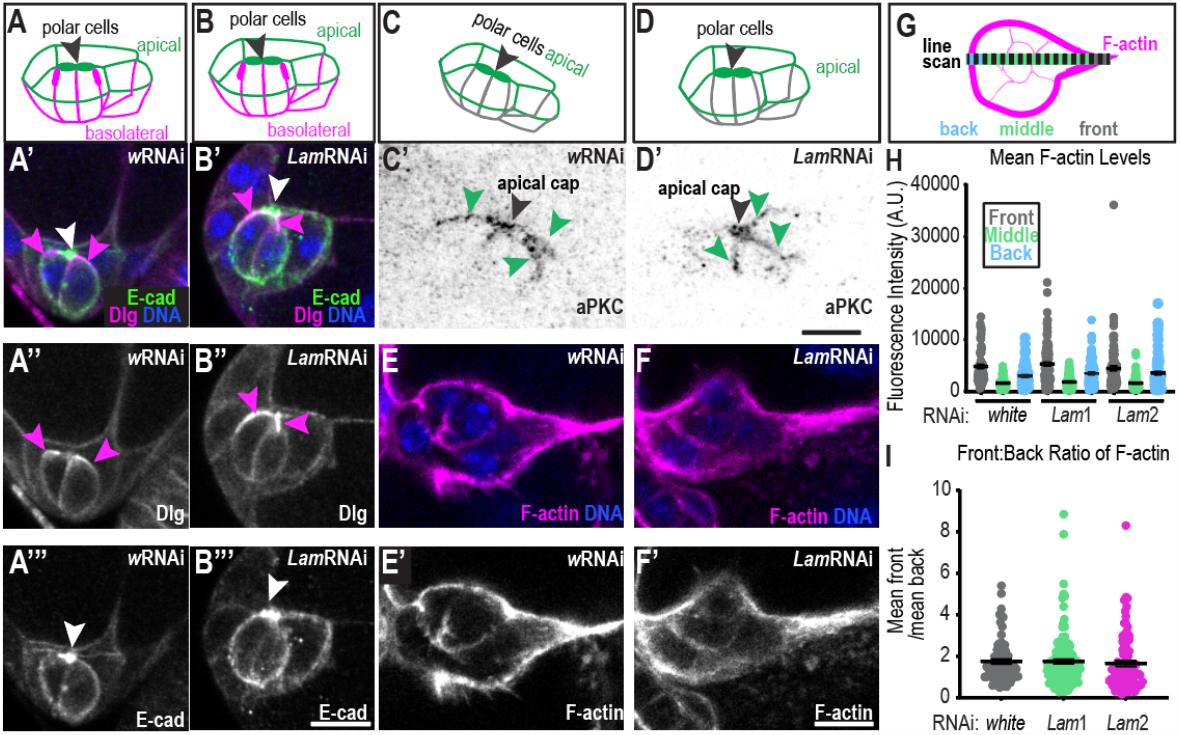
Effects of lamins on cluster polarity. A-B. Schematics of polarity orientation and merged images of control (A’) and LamRNAi (B’) clusters marked with E-cadherin to mark the apical surfaces, Dlg to mark basolateral surfaces, and Hoecsht to mark DNA. A’’-B’’ Dlg greyscale image A’’’-B’’’ E-cadherin greyscale image. White arrowhead: polar cell apical cap, magenta arrowheads: lateral Dlg. C-D. Schematic and and inverted greyscale images (C’-D’) of clusters are stained for aPKC as an apical marker. Black arrowhead: polar cell apical cap, green arrowheads: apical border cell surface. E-F. Images of border cells stained with Phalloidin to mark F-actin and Hoechst to mark DNA. E’-F’ Greyscale images of F-actin. G. Schematic of line scale acquired across cluster. The binning of regions were for Front: 0-10% length, Middle: 11-89% length, and Back: 90-100% length. H. Individual and mean +/-SEM F-actin levels in indicated clusters and position. Statistical test: Brown Forsythe and Welch followed by Dunnett’s Multiple Comparison, means for front, back or middle F-actin are all not significantly different from LamRNAis for relative position, p>0.9. I. Individual and mean +/- SEM ratios of the front: back intensity for each cluster. Statistical test: Kruskal Wallis test, wRNA versus Lam1RNAi: p>0.9, wRNAi versus Lam2RNAi: p=0.24. All KD conditions in figure: 3 days at 29 ºC. Scale bars: 10 ***μ***m. Genotypes and experimental replicates reported in Table 2.

**Figure S5.**
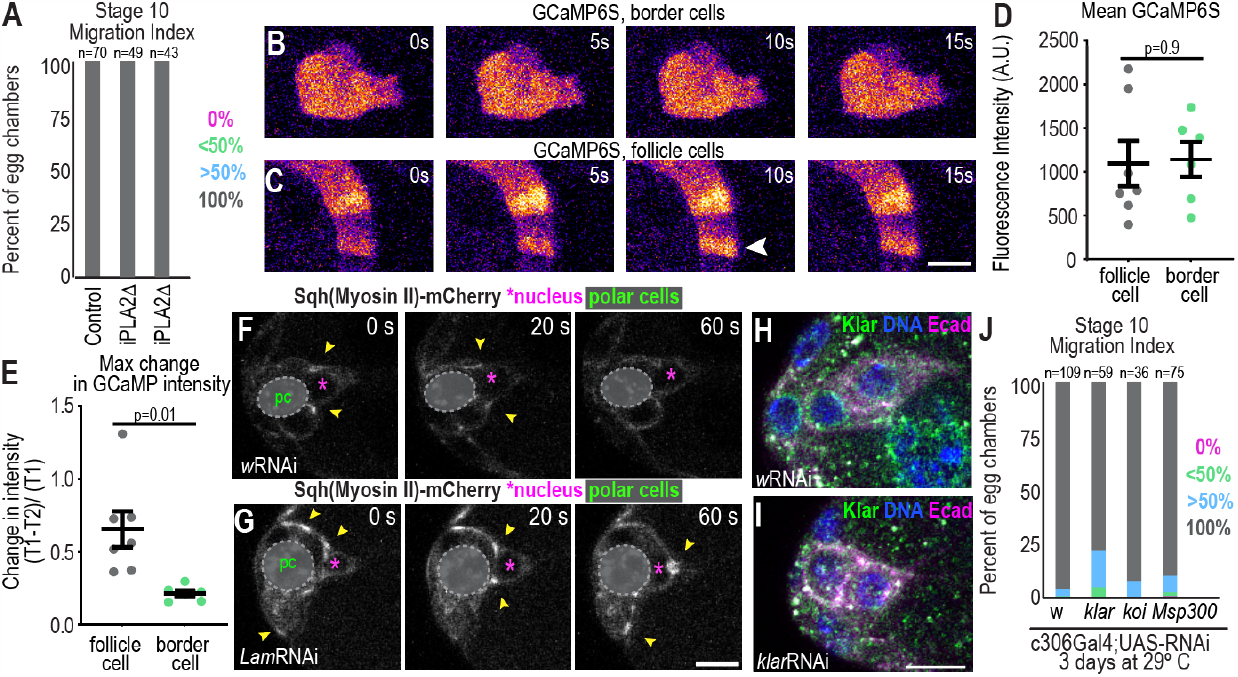
Border cells do not require the LINC complex for migration and lack calcium dynamics but have rapid flashes of myosin around nuclei. A. Migration Indices for the indicated conditions. iPLA2 mutants are iPLA2-VIA[Delta174] and iPLA2-VIA[Delta192], which are both predicted to excise the start codon and have been validated as null alleles by antibody staining (Lin et al., 2018). N=3 experimental replicates. B-C Select images of border cells (B) or follicle cells (C) expressing UAS-GCaMP6S driven by c306Gal4. D. Plot showing the individual and mean +/SEM fluorescent intensity of GCAMP6s. An unpaired t-test was performed. E. Plot of the maximum change in border cells and follicle cells with flashes. Individual (dots) and mean +/- SEM (bars) are shown. A Welch’s t test was performed. F-G Example images of Sqh-mCherry in control (F) and Lam-depleted clusters (G) after 3 day incubation at 29 ºC. H-I. Example images of border cell clusters stained for Klar and indicated markers in control (H) and KlarRNAi (I). J. Migration index for indicated conditions. Fisher’s exact test with Bonferroni correction to compare % with complete migration(wRNAi versus KlarRNAi: p=0.003, wRNAi versus KoiRNAi: p>0.9, and wRNAi versus Msp300, p>0.9). Genotypes and experimental replicates reported in Table 2.

## Supplemental Videos

**Video 1. Border cell cluster undergoing delamination**. Single z-slices from a stage 9 egg chamber expressing fruitlessGal4; UAS-GFP-moesin actin-binding domain (border cells, green) UAS-dsRED.NLS (nuclei, magenta), and incubated with 10 kDa dextran-Alexa 647 (dextrans, cyan). Time interval is 3 minutes. Scale bar: 10 ***μ***m.

**Video 2. The nucleus in the leading cell undergoes rapid changes in shape**. A max projection of a border cell cluster expressing Sqh-mCherry to and GFP.Nuclear localization signal (NLS) at 20 second time intervals. Right: inverted grayscale of GFP.NLS. Scale bar: 10 ***μ***m.

**Video 3. Border cell nuclei undergo dynamic shape changes while polar cells maintain nuclear shape**. Top: Border cell nuclei labeled with UAS-dsRED.NLS and clusters labeled with UAS.GFP-Moesin in lines expressing c306Gal4 (polar cells did not highly express dsRED.NLS). Bottom: Border cell clusters expressing UpdGal4;UAS-dsRED.NLS to label polar cell nuclei and UAS-GFP-moesin to label cells. Time interval is 3 minutes. Scale bar: 10 ***μ***m.

**Video 4. Control border cells maintain a leading protrusion as they delaminate and migrate while Lam-depleted clusters form ectopic protrusions and have undirected movement**. Movies of egg chambers with a control border cells (top, c306Gal4; UAS-wRNAi) or a Lam-depleted cluster (bottom, c306Gal4;UAS-Lam2RNAi) expressing slbo4XPH-EGFP to mark the border cell membranes (green) and UbiHisRFP to mark all nuclei (magenta). Right panel shows slbo4XPH-EGFP channel in grayscale. Time interval is 3 minutes. Scale bar: 10 ***μ***m.

**Video 5. Nuclear stretching and movement corresponds with protrusion widening while Lam-depleted cells do not maintain protrusions**. Timelapse movies of controlRNAi border cells (top, c306Gal4; UAS-wRNAi) or Lam-depleted border cells (bottoms, c306Gal4; UAS-Lam2RNAi) expressing slbo4XPHEGFP to mark the border cells (green) and UbiHisRFP to mark all nuclei (magenta). Time interval is 3 minutes. Right panels show greyscale of UbiHisRFP. Scale bar: 10 ***μ***m.

**Video 6. Photoactivation of GFP-alpha tubulin** Time lapse movie of border cells expressing slboLifeAct RFP and UAS-PA-GFPalphatubulin and photoactivated at a region of interest in a follicle cell (A) or border cell (B). Time interval is 2 seconds. Scale bar: 10 ***μ***m.

**Video 7. Calcium dynamics in border cells**. Time lapse series of egg chambers expressing c306Gal4; UAS-GCaMP6s. Time interval is 5 seconds. Scale bar: 10 ***μ***m.

**Video 8. Myosin II dynamics in control and Lam-depleted clusters** Time lapse movies of border cells expressing Sqh-mCherry for control(left) or Lam-depleted (right) clusters. Polar cells are obscured with a black circle noted at the start of the movie to focus on the border cell cortical flashes. Time interval is 20 seconds. Scale bar: 10 ***μ***m.

**Video 9. Myosin II flashes correspond with nuclear movement**. Time lapse series of border cells expressing Sqh-mCherry and UAS-GFP.NLS driven by c306Gal4. Time interval is 20 seconds. Scale bar: 10 ***μ***m.

**Video 10. Myosin II depletion results in long-lived protrusions and little nuclear movement**. Time lapse series of an egg chamber expressing UbiHisRFP, slbo4XPH-EGFP, c306Gal4;UAS-SqhRNAi after incubation at 29 ºC for 3 days. Scale bar: 10 ***μ***m.

## References

Aranjuez, G., A. Burtscher, K. Sawant, P. Majumder, and J.A. McDonald. 2016. Dynamic myosin activation promotes collective morphology and migration by locally balancing oppositional forces from surrounding tissue. Mol. Biol. Cell. 27:1898–1910. doi:10.1091/mbc.E15-10-0744.

Assaker, G., D. Ramel, S.K. Wculek, M. González-Gaitán, and G. Emery. 2010. Spatial restriction of receptor tyrosine kinase activity through a polarized endocytic cycle controls border cell migration. Proc Natl Acad Sci USA. 107:22558–22563. doi:10.1073/pnas.1010795108.

Beadle, C., M.C. Assanah, P. Monzo, R. Vallee, S.S. Rosenfeld, and P. Canoll. 2008. The role of myosin II in glioma invasion of the brain. Mol. Biol. Cell. 19:3357–3368. doi:10.1091/mbc.E08-03-0319.

Belyaeva, V., S. Wachner, A. Gyoergy, S. Emtenani, I. Gridchyn, M. Akhmanova, M. Linder, M. Roblek, M. Sibilia, and D. Siekhaus. 2022. Fos regulates macrophage infiltration against surrounding tissue resistance by a cortical actin-based mechanism in Drosophila. PLoS Biol. 20:e3001494. doi:10.1371/journal.pbio.3001494.

Ben-David, G., E. Miller, and J. Steinhauer. 2015. Drosophila spermatid individualization is sensitive to temperature and fatty acid metabolism. Spermatogenesis. 5:e1006089. doi:10.1080/21565562.2015.1006089.

Bossie, C.A., and M.M. Sanders. 1993. A cDNA from Drosophila melanogaster encodes a lamin C-like intermediate filament protein. J. Cell Sci. 104 (Pt 4):1263–1272. doi:10.1242/jcs.104.4.1263.

Cai, D., S.-C. Chen, M. Prasad, L. He, X. Wang, V. Choesmel-Cadamuro, J.K. Sawyer, G. Danuser, and D.J. Montell. 2014. Mechanical feedback through E-cadherin promotes direction sensing during collective cell migration. Cell. 157:1146–1159. doi:10.1016/j.cell.2014.03.045.

Calero-Cuenca, F.J., C.S. Janota, and E.R. Gomes. 2018. Dealing with the nucleus during cell migration. Curr. Opin. Cell Biol. 50:35–41. doi:10.1016/j.ceb.2018.01.014.

Campanale, J.P., J.A. Mondo, and D.J. Montell. 2022. A Scribble/Cdep/Rac pathway controls follower-cell crawling and cluster cohesion during collective border-cell migration. Dev. Cell. 57:2483-2496.e4. doi:10.1016/j.devcel.2022.10.004.

Chen, N.Y., Y. Yang, T.A. Weston, J.N. Belling, P. Heizer, Y. Tu, P. Kim, L. Edillo, S.J. Jonas, P.S. Weiss, L.G. Fong, and S.G. Young. 2019. An absence of lamin B1 in migrating neurons causes nuclear membrane ruptures and cell death. Proc Natl Acad Sci USA. 116:25870–25879. doi:10.1073/pnas.1917225116.

Collins, M.A., T.R. Mandigo, J.M. Camuglia, G.A. Vazquez, A.J. Anderson, C.H. Hudson, J.L. Hanron, and E.S. Folker. 2017. Emery-Dreifuss muscular dystrophy-linked genes and centronuclear myopathy-linked genes regulate myonuclear movement by distinct mechanisms. Mol. Biol. Cell. 28:2303–2317. doi:10.1091/mbc.E16-10-0721.

Crisp, M., Q. Liu, K. Roux, J.B. Rattner, C. Shanahan, B. Burke, P.D. Stahl, and D. Hodzic. 2006. Coupling of the nucleus and cytoplasm: role of the LINC complex. J. Cell Biol. 172:41–53. doi:10.1083/jcb.200509124.

Dai, W., X. Guo, Y. Cao, J.A. Mondo, J.P. Campanale, B.J. Montell, H. Burrous, S. Streichan, N. Gov, W.-J. Rappel, and D.J. Montell. 2020. Tissue topography steers migrating Drosophila border cells. Science. 370:987–990. doi:10.1126/science.aaz4741.

Davidson, P.M., C. Denais, M.C. Bakshi, and J. Lammerding. 2014. Nuclear deformability constitutes a rate-limiting step during cell migration in 3-D environments. Cell. Mol. Bioeng. 7:293–306. doi:10.1007/s12195-014-0342-y.

Davidson, P.M., and J. Lammerding. 2014. Broken nuclei--lamins, nuclear mechanics, and disease. Trends Cell Biol. 24:247–256. doi:10.1016/j.tcb.2013.11.004.

Denais, C.M., R.M. Gilbert, P. Isermann, A.L. McGregor, M. te Lindert, B. Weigelin, P.M. Davidson, P. Friedl, K. Wolf, and J. Lammerding. 2016. Nuclear envelope rupture and repair during cancer cell migration. Science. 352:353–358. doi:10.1126/science.aad7297.

Duchek, P., K. Somogyi, G. Jékely, S. Beccari, and P. Rørth. 2001. Guidance of cell migration by the Drosophila PDGF/VEGF receptor. Cell. 107:17–26. doi:10.1016/s0092-8674(01)00502-5.

Ferrera, D., C. Canale, R. Marotta, N. Mazzaro, M. Gritti, M. Mazzanti, S. Capellari, P. Cortelli, and L. Gasparini. 2014. Lamin B1 overexpression increases nuclear rigidity in autosomal dominant leukodystrophy fibroblasts. FASEB J. 28:3906–3918. doi:10.1096/fj.13-247635.

Friedl, P., and D. Gilmour. 2009. Collective cell migration in morphogenesis, regeneration and cancer. Nat. Rev. Mol. Cell Biol. 10:445–457. doi:10.1038/nrm2720.

Friedl, P., K. Wolf, and J. Lammerding. 2011. Nuclear mechanics during cell migration. Curr. Opin. Cell Biol. 23:55–64. doi:10.1016/j.ceb.2010.10.015.

Graham, D.M., T. Andersen, L. Sharek, G. Uzer, K. Rothenberg, B.D. Hoffman, J. Rubin, M. Balland, J.E. Bear, and K. Burridge. 2018. Enucleated cells reveal differential roles of the nucleus in cell migration, polarity, and mechanotransduction. J. Cell Biol. 217:895–914. doi:10.1083/jcb.201706097.

Hampoelz, B., M.-T. Mackmull, P. Machado, P. Ronchi, K.H. Bui, N. Schieber, R. Santarella-Mellwig, A. Necakov, A. Andrés-Pons, J.M. Philippe, T. Lecuit, Y. Schwab, and M. Beck. 2016. Pre-assembled Nuclear Pores Insert into the Nuclear Envelope during Early Development. Cell. 166:664–678. doi:10.1016/j.cell.2016.06.015.

Harada, T., J. Swift, J. Irianto, J.-W. Shin, K.R. Spinler, A. Athirasala, R. Diegmiller, P.C.D.P. Dingal, I.L. Ivanovska, and D.E. Discher. 2014. Nuclear lamin stiffness is a barrier to 3D migration, but softness can limit survival. J. Cell Biol. 204:669–682. doi:10.1083/jcb.201308029.

Hetzer, M.W. 2010. The nuclear envelope. Cold Spring Harb. Perspect. Biol. 2:a000539. doi:10.1101/cshperspect.a000539.

Hoffmann, K., C.K. Dreger, A.L. Olins, D.E. Olins, L.D. Shultz, B. Lucke, H. Karl, R. Kaps, D. Müller, A. Vayá, J. Aznar, R.E. Ware, N. Sotelo Cruz, T.H. Lindner, H. Herrmann, A. Reis, and K. Sperling. 2002. Mutations in the gene encoding the lamin B receptor produce an altered nuclear morphology in granulocytes (Pelger-Huët anomaly). Nat. Genet. 31:410–414. doi:10.1038/ng925.

Hutchison, C.J. 2014. B-type lamins in health and disease. Semin. Cell Dev. Biol. 29:158–163. doi:10.1016/j.semcdb.2013.12.012.

Irianto, J., Y. Xia, C.R. Pfeifer, A. Athirasala, J. Ji, C. Alvey, M. Tewari, R.R. Bennett, S.M. Harding, A.J. Liu, R.A. Greenberg, and D.E. Discher. 2017. DNA Damage Follows Repair Factor Depletion and Portends Genome Variation in Cancer Cells after Pore Migration. Curr. Biol. 27:210–223. doi:10.1016/j.cub.2016.11.049.

Jahed, Z., and M.R. Mofrad. 2019. The nucleus feels the force, LINCed in or not! Curr. Opin. Cell Biol. 58:114–119. doi:10.1016/j.ceb.2019.02.012.

Johnston, I.D., D.K. McCluskey, C.K.L. Tan, and M.C. Tracey. 2014. Mechanical characterization of bulk Sylgard 184 for microfluidics and microengineering. J. Micromech. Microeng. 24:035017. doi:10.1088/0960-1317/24/3/035017.

Jung, H.-J., C. Coffinier, Y. Choe, A.P. Beigneux, B.S.J. Davies, S.H. Yang, R.H. Barnes, J. Hong, T. Sun, S.J. Pleasure, S.G. Young, and L.G. Fong. 2012. Regulation of prelamin A but not lamin C by miR-9, a brain-specific microRNA. Proc Natl Acad Sci USA. 109:E423–31. doi:10.1073/pnas.1111780109.

Kalukula, Y., A.D. Stephens, J. Lammerding, and S. Gabriele. 2022. Mechanics and functional consequences of nuclear deformations. Nat. Rev. Mol. Cell Biol. 23:583–602. doi:10.1038/s41580-022-00480-z.

Lamb, M.C., C.P. Kaluarachchi, T.I. Lansakara, S.Q. Mellentine, Y. Lan, A.V. Tivanski, and T.L. Tootle. 2021. Fascin limits Myosin activity within Drosophila border cells to control substrate stiffness and promote migration. eLife. 10. doi:10.7554/eLife.69836.

Lammerding, J., P.C. Schulze, T. Takahashi, S. Kozlov, T. Sullivan, R.D. Kamm, C.L. Stewart, and R.T. Lee. 2004. Lamin A/C deficiency causes defective nuclear mechanics and mechanotransduction. J. Clin. Invest. 113:370–378. doi:10.1172/JCI19670.

Lee, H.-P., F. Alisafaei, K. Adebawale, J. Chang, V.B. Shenoy, and O. Chaudhuri. 2021. The nuclear piston activates mechanosensitive ion channels to generate cell migration paths in confining microenvironments. Sci. Adv. 7. doi:10.1126/sciadv.abd4058.

Lee, J.S.H., C.M. Hale, P. Panorchan, S.B. Khatau, J.P. George, Y. Tseng, C.L. Stewart, D. Hodzic, and D. Wirtz. 2007. Nuclear lamin A/C deficiency induces defects in cell mechanics, polarization, and migration. Biophys. J. 93:2542–2552. doi:10.1529/biophysj.106.102426.

Lenz-Böhme, B., J. Wismar, S. Fuchs, R. Reifegerste, E. Buchner, H. Betz, and B. Schmitt. 1997. Insertional mutation of the Drosophila nuclear lamin Dm0 gene results in defective nuclear envelopes, clustering of nuclear pore complexes, and accumulation of annulate lamellae. J. Cell Biol. 137:1001–1016. doi:10.1083/jcb.137.5.1001.

Lin, G., P.-T. Lee, K. Chen, D. Mao, K.L. Tan, Z. Zuo, W.-W. Lin, L. Wang, and H.J. Bellen. 2018. Phospholipase PLA2G6, a Parkinsonism-Associated Gene, Affects Vps26 and Vps35, Retromer Function, and Ceramide Levels, Similar to α-Synuclein Gain. Cell Metab. 28:605-618.e6. doi:10.1016/j.cmet.2018.05.019.

Lomakin, A.J., C.J. Cattin, D. Cuvelier, Z. Alraies, M. Molina, G.P.F. Nader, N. Srivastava, P.J. Sáez, J.M. Garcia-Arcos, I.Y. Zhitnyak, A. Bhargava, M.K. Driscoll, E.S. Welf, R. Fiolka, R.J. Petrie, N.S. De Silva, J.M. González-Granado, N. Manel, A.M. Lennon-Duménil, D.J. Müller, and M. Piel. 2020. The nucleus acts as a ruler tailoring cell responses to spatial constraints. Science. 370. doi:10.1126/science.aba2894.

Luo, J., P. Zhou, X. Guo, D. Wang, and J. Chen. 2019. The polarity protein Dlg5 regulates collective cell migration during Drosophila oogenesis. PLoS ONE. 14:e0226061. doi:10.1371/journal.pone.0226061.

Majumder, P., G. Aranjuez, J. Amick, and J.A. McDonald. 2012. Par-1 controls myosin-II activity through myosin phosphatase to regulate border cell migration. Curr. Biol. 22:363–372. doi:10.1016/j.cub.2012.01.037.

McDonald, J.A., A. Khodyakova, G. Aranjuez, C. Dudley, and D.J. Montell. 2008. PAR-1 kinase regulates epithelial detachment and directional protrusion of migrating border cells. Curr. Biol. 18:1659–1667. doi:10.1016/j.cub.2008.09.041.

McGregor, A.L., C.-R. Hsia, and J. Lammerding. 2016. Squish and squeeze-the nucleus as a physical barrier during migration in confined environments. Curr. Opin. Cell Biol. 40:32–40. doi:10.1016/j.ceb.2016.01.011.

Mishra, A.K., J.P. Campanale, J.A. Mondo, and D.J. Montell. 2019a. Cell interactions in collective cell migration. Development. 146. doi:10.1242/dev.172056.

Mishra, A.K., J.A. Mondo, J.P. Campanale, and D.J. Montell. 2019b. Coordination of protrusion dynamics within and between collectively migrating border cells by myosin II. Mol. Biol. Cell. 30:2490–2502. doi:10.1091/mbc.E19-02-0124.

Montell, D.J., W.H. Yoon, and M. Starz-Gaiano. 2012. Group choreography: mechanisms orchestrating the collective movement of border cells. Nat. Rev. Mol. Cell Biol. 13:631–645. doi:10.1038/nrm3433.

Murphy, A.M., and D.J. Montell. 1996. Cell type-specific roles for Cdc42, Rac, and RhoL in Drosophila oogenesis. J. Cell Biol. 133:617–630. doi:10.1083/jcb.133.3.617.

Norden, C., S. Young, B.A. Link, and W.A. Harris. 2009. Actomyosin is the main driver of interkinetic nuclear migration in the retina. Cell. 138:1195–1208. doi:10.1016/j.cell.2009.06.032.

Perillo, M., and E.S. Folker. 2018. Specialized Positioning of Myonuclei Near Cell-Cell Junctions. Front. Physiol. 9:1531. doi:10.3389/fphys.2018.01531.

Petrie, R.J., H. Koo, and K.M. Yamada. 2014. Generation of compartmentalized pressure by a nuclear piston governs cell motility in a 3D matrix. Science. 345:1062–1065. doi:10.1126/science.1256965.

Petrovsky, R., and J. Großhans. 2018. Expression of lamina proteins Lamin and Kugelkern suppresses stem cell proliferation. Nucleus. 9:104–118. doi:10.1080/19491034.2017.1412028.

Pinheiro, E.M., and D.J. Montell. 2004. Requirement for Par-6 and Bazooka in Drosophila border cell migration. Development. 131:5243–5251. doi:10.1242/dev.01412.

Prüfert, K., A. Vogel, and G. Krohne. 2004. The lamin CxxM motif promotes nuclear membrane growth. J. Cell Sci. 117:6105–6116. doi:10.1242/jcs.01532.

Raab, M., M. Gentili, H. de Belly, H.R. Thiam, P. Vargas, A.J. Jimenez, F. Lautenschlaeger, R. Voituriez, A.M. Lennon-Duménil, N. Manel, and M. Piel. 2016. ESCRT III repairs nuclear envelope ruptures during cell migration to limit DNA damage and cell death. Science. 352:359–362. doi:10.1126/science.aad7611.

Renkawitz, J., A. Kopf, J. Stopp, I. de Vries, M.K. Driscoll, J. Merrin, R. Hauschild, E.S. Welf, G. Danuser, R. Fiolka, and M. Sixt. 2019. Nuclear positioning facilitates amoeboid migration along the path of least resistance. Nature. 568:546–550. doi:10.1038/s41586-019-1087-5.

Reymond, N. B.B. d’Água, and A.J. Ridley. 2013. Crossing the endothelial barrier during metastasis. Nat. Rev. Cancer. 13:858–870. doi:10.1038/nrc3628.

Rørth, P. 2009. Collective cell migration. Annu. Rev. Cell Dev. Biol. 25:407–429. doi:10.1146/annurev.cellbio.042308.113231.

Rowat, A.C., D.E. Jaalouk, M. Zwerger, W.L. Ung, I.A. Eydelnant, D.E. Olins, A.L. Olins, H. Herrmann, D.A. Weitz, and J. Lammerding. 2013. Nuclear envelope composition determines the ability of neutrophil-type cells to passage through micron-scale constrictions. J. Biol. Chem. 288:8610–8618. doi:10.1074/jbc.M112.441535.

Sahu, A., R. Ghosh, G. Deshpande, and M. Prasad. 2017. A Gap Junction Protein, Inx2, Modulates Calcium Flux to Specify Border Cell Fate during Drosophila oogenesis. PLoS Genet. 13:e1006542. doi:10.1371/journal.pgen.1006542.

Shin, J.-W., K.R. Spinler, J. Swift, J.A. Chasis, N. Mohandas, and D.E. Discher. 2013. Lamins regulate cell trafficking and lineage maturation of adult human hematopoietic cells. Proc Natl Acad Sci USA. 110:18892–18897. doi:10.1073/pnas.1304996110.

Silver, D.L., E.R. Geisbrecht, and D.J. Montell. 2005. Requirement for JAK/STAT signaling throughout border cell migration in Drosophila. Development. 132:3483–3492. doi:10.1242/dev.01910.

Silver, D.L., and D.J. Montell. 2001. Paracrine signaling through the JAK/STAT pathway activates invasive behavior of ovarian epithelial cells in Drosophila. Cell. 107:831–841. doi:10.1016/s0092-8674(01)00607-9.

Stoletov, K., H. Kato, E. Zardouzian, J. Kelber, J. Yang, S. Shattil, and R. Klemke. 2010. Visualizing extravasation dynamics of metastatic tumor cells. J. Cell Sci. 123:2332–2341. doi:10.1242/jcs.069443.

Swift, J., I.L. Ivanovska, A. Buxboim, T. Harada, P.C.D.P. Dingal, J. Pinter, J.D. Pajerowski, K.R. Spinler, J.-W. Shin, M. Tewari, F. Rehfeldt, D.W. Speicher, and D.E. Discher. 2013. Nuclear lamin-A scales with tissue stiffness and enhances matrix-directed differentiation. Science. 341:1240104. doi:10.1126/science.1240104.

Thiam, H.-R., P. Vargas, N. Carpi, C.L. Crespo, M. Raab, E. Terriac, M.C. King, J. Jacobelli, A.S. Alberts, T. Stradal, A.-M. Lennon-Dumenil, and M. Piel. 2016. Perinuclear Arp2/3-driven actin polymerization enables nuclear deformation to facilitate cell migration through complex environments. Nat. Commun. 7:10997. doi:10.1038/ncomms10997.

Vargas, J.D., E.M. Hatch, D.J. Anderson, and M.W. Hetzer. 2012. Transient nuclear envelope rupturing during interphase in human cancer cells. Nucleus. 3:88–100. doi:10.4161/nucl.18954.

Venturini, V., F. Pezzano, F. Català Castro, H.-M. Häkkinen, S. Jiménez-Delgado, M. Colomer-Rosell, M. Marro, Q. Tolosa-Ramon, S. Paz-López, M.A. Valverde, J. Weghuber, P. Loza-Alvarez, M. Krieg, S. Wieser, and V. Ruprecht. 2020. The nucleus measures shape changes for cellular proprioception to control dynamic cell behavior. Science. 370. doi:10.1126/science.aba2644.

Wang, H., Z. Qiu, Z. Xu, S.J. Chen, J. Luo, X. Wang, and J. Chen. 2018. aPKC is a key polarity determinant in coordinating the function of three distinct cell polarities during collective migration. Development. 145. doi:10.1242/dev.158444.

Wang, X., J.C. Adam, and D. Montell. 2007. Spatially localized Kuzbanian required for specific activation of Notch during border cell migration. Dev. Biol. 301:532–540. doi:10.1016/j.ydbio.2006.08.031.

Wintner, O., N. Hirsch-Attas, M. Schlossberg, F. Brofman, R. Friedman, M. Kupervaser, D. Kitsberg, and A. Buxboim. 2020. A unified linear viscoelastic model of the cell nucleus defines the mechanical contributions of lamins and chromatin. Adv Sci (Weinh). 7:1901222. doi:10.1002/advs.201901222.

Wolf, K., M. Te Lindert, M. Krause, S. Alexander, J. Te Riet, A.L. Willis, R.M. Hoffman, C.G. Figdor, S.J. Weiss, and P. Friedl. 2013. Physical limits of cell migration: control by ECM space and nuclear deformation and tuning by proteolysis and traction force. J. Cell Biol. 201:1069–1084. doi:10.1083/jcb.201210152.

Wolf, K., Y.I. Wu, Y. Liu, J. Geiger, E. Tam, C. Overall, M.S. Stack, and P. Friedl. 2007. Multi-step pericellular proteolysis controls the transition from individual to collective cancer cell invasion. Nat. Cell Biol. 9:893–904. doi:10.1038/ncb1616.

Wu, J.-W., C.-W. Wang, R.-Y. Chen, L.-Y. Hung, Y.-C. Tsai, Y.-T. Chan, Y.-C. Chang, and A.C.-C. Jang. 2022. Spatiotemporal gating of Stat nuclear influx by Drosophila Npas4 in collective cell migration. Sci. Adv. 8:eabm2411. doi:10.1126/sciadv.abm2411.

Wyckoff, J.B., J.G. Jones, J.S. Condeelis, and J.E. Segall. 2000. A critical step in metastasis: in vivo analysis of intravasation at the primary tumor. Cancer Res. 60:2504–2511.

Xie, W., A. Chojnowski, T. Boudier, J.S.Y. Lim, S. Ahmed, Z. Ser, C. Stewart, and B. Burke. 2016. A-type Lamins Form Distinct Filamentous Networks with Differential Nuclear Pore Complex Associations. Curr. Biol. 26:2651–2658. doi:10.1016/j.cub.2016.07.049.

Yamada, K.M., and M. Sixt. 2019. Mechanisms of 3D cell migration. Nat. Rev. Mol. Cell Biol. 20:738–752. doi:10.1038/s41580-019-0172-9.

Yamauchi, K., M. Yang, P. Jiang, N. Yamamoto, M. Xu, Y. Amoh, K. Tsuji, M. Bouvet, H. Tsuchiya, K. Tomita, A.R. Moossa, and R.M. Hoffman. 2005. Real-time in vivo dual-color imaging of intracapillary cancer cell and nucleus deformation and migration. Cancer Res. 65:4246–4252. doi:10.1158/0008-5472.CAN-05-0069.

Zacharioudaki, E., and S.J. Bray. 2014. Tools and methods for studying Notch signaling in Drosophila melanogaster. Methods. 68:173–182. doi:10.1016/j.ymeth.2014.03.029

